# IqgD is a Rac1-interacting IQGAP required for efficient growth of *Dictyostelium discoideum* on bacterial lawns

**DOI:** 10.64898/2026.02.02.703220

**Authors:** Anja Čizmar, Darija Putar, Marija Šimić, Jonas Scholz, Maja Marinović, Lucija Horvat, Mihaela Matovina, Igor Weber, Jan Faix, Vedrana Filić

## Abstract

IQGAPs are large multidomain scaffold proteins that interact with the Rho family GTPases Cdc42 and Rac1, functioning both as their effectors and as regulators by stabilizing their active GTP-bound state. In this study, we analyzed the function of IqgD, an IQGAP-related protein from the professional phagocyte *Dictyostelium discoideum*. IqgD contains a calponin homology domain (CHD), a GAP-related domain (GRD), and a RasGAP C-terminal (RGCt) domain. We show that the CHD is essential for F-actin binding and cortical localization, whereas the GRD and RGCt domains mediate interactions with Rac1 GTPases and the actin-bundling proteins cortexillins. Moreover, similar to mammalian IQGAPs, IqgD maintains Rac1 in its active conformation. IqgD is enriched in macropinocytic and phagocytic cups and co-localizes with F-actin and active Rac1 in the ring-like structure that forms around surface-bound particles at the cell bottom. Loss of IqgD results in markedly reduced growth on bacterial lawns and significantly smaller cell size. While mutant cells internalize bacteria from suspension as efficiently as wild-type cells, they display a strong defect in phagocytosis of surface-bound particles, accompanied by decreased adhesion to the cell substrate. Together, our data show that although IqgD localizes to macroendocytic cups, it is dispensable for macropinocytosis and phagocytosis of suspended particles. Instead, IqgD is specifically required for efficient phagocytosis of surface-bound particles, likely by facilitating robust F-actin polymerization at the cell bottom to generate the force necessary for detachment of surface-bound bacteria.

**Significance Statement:** Phagocytosis of surface-bound microbes is essential for host defense and environmental feeding strategies, yet its underlying mechanisms remain poorly understood. We identify the IQGAP-related protein IqgD in *D. discoideum* as a key factor required for efficient uptake of bacteria attached to solid surfaces. IqgD localizes to an F-actin- and Rac1-rich circular structure analogous to the phagocytic adhesion ring (PAR) recently described in mammalian macrophages, suggesting that this mode of force-driven particle detachment is evolutionarily conserved. Our findings provide mechanistic insight into substrate-dependent phagocytosis and establish IqgD as a central regulator of this process.

## Introduction

IQ motif-containing GTPase-activating proteins (IQGAPs) constitute an evolutionarily conserved protein family in eukaryotes. Most vertebrates express three isoforms with similar domain architecture: an N-terminal calponin homology domain (CHD), followed by a coiled-coil repeat region (CC), a tryptophan-containing proline-rich motif-binding region (WW), an IQ domain containing several IQ motifs, a GTPase-activating protein (GAP)-related domain (GRD), a RasGAP C-terminal (RGCt) domain, and an extreme C-terminal (CT) domain (1, 2). These multidomain proteins act as scaffolds that modulate various signaling pathways by bringing their components into close proximity. For example, the best-studied IQGAP, human IQGAP1, promotes the mitogen-activated protein kinase (MAPK) signaling pathway by directly binding B-Raf, C-Raf, MEK1/2, and ERK1/2 (3–5). IQGAP1 also facilitates PI3K/Akt signaling by binding three phosphoinositide kinases that sequentially generate PtdIns(3,4,5)P_3_ from phosphatidylinositol, as well as Akt and PDK1, which are recruited to PtdIns(3,4,5)P_3_ (6). IQGAPs interact with various cell surface receptors and serve as scaffolds for their signaling components, modulating their activation and signaling (7). Last but not least, IQGAPs are important regulators of the actin cytoskeleton (8).

Although IQGAPs harbor the GRD, which is structurally very similar to the RasGAP domain, their catalytic activity to promote GTP hydrolysis on Ras GTPases appears to have been lost early in evolution due to mutations of key amino acid residues (9–11). Furthermore, IQGAPs do not bind the canonical GTPases of the Ras family (12). Instead, they interact with the Rho family GTPases, Cdc42 and Rac1, and stabilize their active forms (13–17). Overexpression and silencing of IQGAP1 leads to increased and decreased levels of active Cdc42, respectively (18–20). Binding of Cdc42 to IQGAP1 requires GRD, but additional regions, particularly in the C-terminal part of IQGAP1, contribute to GTPase binding (13, 18–20). More recently, a multistep binding mechanism has been proposed (21, 22). The authors suggested that RGCt is the central domain that initially associates with the switch regions of GTP-loaded Cdc42 and Rac1, while GRD subsequently mediates low-affinity, partially nucleotide-dependent binding outside the switch regions. The CT domain binds with extremely low affinity outside the switch regions and most likely stabilizes the interaction between IQGAP1 and Cdc42.

IQGAPs influence the activity of Rho family GTPases by maintaining their activated state, but they also act as their effectors (23). IQGAPs bind microfilaments directly via the CHD and cross-link them into loosely organized bundles (24–26). This activity is dependent on their oligomerization, which is enhanced by active Cdc42 (25). Furthermore, IQGAP1 directly influences actin dynamics as a barbed end capper that transiently arrests filament growth and prevents depolymerization from the barbed end (26, 27). In addition, IQGAP1 interacts with N-WASP, although there are conflicting data on whether it activates N-WASP and thereby stimulates Arp2/3-mediated nucleation of branched actin filaments (27–30). Interestingly, recent work has shown that IQGAP1 regulates filament growth by displacing barbed end binding proteins such as formin Dia1 or the complex of Dia1 and the capping factor CP, which determines the polymerization fate of a single filament (31, 32). Owing to these functions, IQGAPs are involved in fundamental cellular processes, including cell migration, cytokinesis and endocytosis (33).

The protist *Dictyostelium discoideum* expresses four proteins with adjacent GRD and RGCt domains present in all IQGAPs: DGAP1, GAPA, IqgC, and IqgD (34). However, DGAP1, GAPA, and IqgC lack the N-terminal domains found in IQGAPs from higher eukaryotes. Therefore, they are unable to interact directly with actin filaments. Interestingly, DGAP1 and GAPA interact with actin-binding proteins, the cortexillins, which are found only in amebozoa, and bridge the gap between these truncated IQGAPs and F-actin (35). The association of DGAP1 or GAPA with a cortexillin dimer requires the binding of active Rac1 to DGAP1 or GAPA. The Rho family of GTPases in *D. discoideum* comprises 20 Rac GTPases, 6 of which belong to the Rac subfamily (Rac1A, Rac1B, Rac1C, RacF1, RacF2, and RacB) (36). Rac1A, Rac1B, and Rac1C are *D. discoideum* orthologs of human Rac1. They are more than 90% identical in amino acid sequence and have an identical effector-binding domain. The above-mentioned tetrameric complex of active Rac1, DGAP1 or GAPA, and a cortexillin dimer forms in the posterior cortex of interphase cells and in the cleavage furrow of dividing cells, and is important for efficient migration and cytokinesis (35, 37, 38). Functional characterization of IqgC revealed that, in contrast to DGAP1 and GAPA, IqgC exhibits RasGAP activity and is involved in the regulation of large-scale endocytosis (39, 40). This is consistent with phylogenetic analysis, which places IqgC closer to the Gap1 family of RasGAPs from fungi than to IQGAPs from *Dictyostelium* (11, 41). IqgD is the only IQGAP-related protein from *D. discoideum* that contains an N-terminal CHD and is significantly larger than DGAP1, GAPA, and IqgC. Nevertheless, apart from the CHD, GRD, and RGCt, it does not contain any other domain characteristic of IQGAPs. Here, we show that IqgD binds to Rac1 GTPases, F-actin, and cortexillins. Binding to F-actin is mediated via the CHD, whereas interactions with Rac1 and cortexillin I/II require the GRD and RGCt domains of IqgD. Furthermore, we show that IqgD maintains Rac1 GTPase in an active state and that Rac1-bound IqgD cannot interact with cortexillin. IqgD is localized in cortical structures and is strongly enriched in macropinocytic and phagocytic cups. It also accumulates in circular structures at the cell bottom that form around surface-bound particles. Comprehensive phenotypic analyses of cells lacking IqgD showed that it is required for efficient growth on bacterial lawns. Further assays showed that IqgD-deficient cells display impaired substrate adhesion and a reduced ability to consume particles attached to the surface, which likely prevents them from effectively grazing bacterial lawns.

## Results

### IqgD localizes to F-actin-rich structures

To investigate the role of IqgD during growth, we examined the subcellular localization of fluorescently labeled IqgD during random movement, phagocytosis, and cytokinesis of vegetative wild-type cells (Fig. 1). IqgD was fused at both the N- and C-termini to yellow fluorescent protein (YFP), and both fusion proteins localized throughout the cell cortex and were strongly enriched in macropinocytic and phagocytic cups (Fig. 1A-C and Fig. S1A). While IqgD appeared evenly distributed on developing macropinosomes, it was markedly enriched at the base of forming phagocytic cups. As phagocytosis progressed, IqgD shifted from the base to distal regions of the cup and accumulated strongly at the site of cup closure. During cytokinesis, IqgD was approximately evenly distributed throughout the cortex (Fig. 1D). The localization of IqgD during growth closely resembled the distribution of F-actin in vegetative cells. Consistent with this, co-expression of IqgD-YFP (Fig. S1A) or YFP-IqgD (Fig. S1B) with the F-actin marker Lifeact-mRFP (42) revealed almost complete colocalization at the cell cortex and within macropinocytic and phagocytic cups. Since YFP-IqgD labeled the cell cortex more strongly than IqgD-YFP, all subsequent localization experiments were performed using the N-terminally tagged construct.

**Figure 1.**
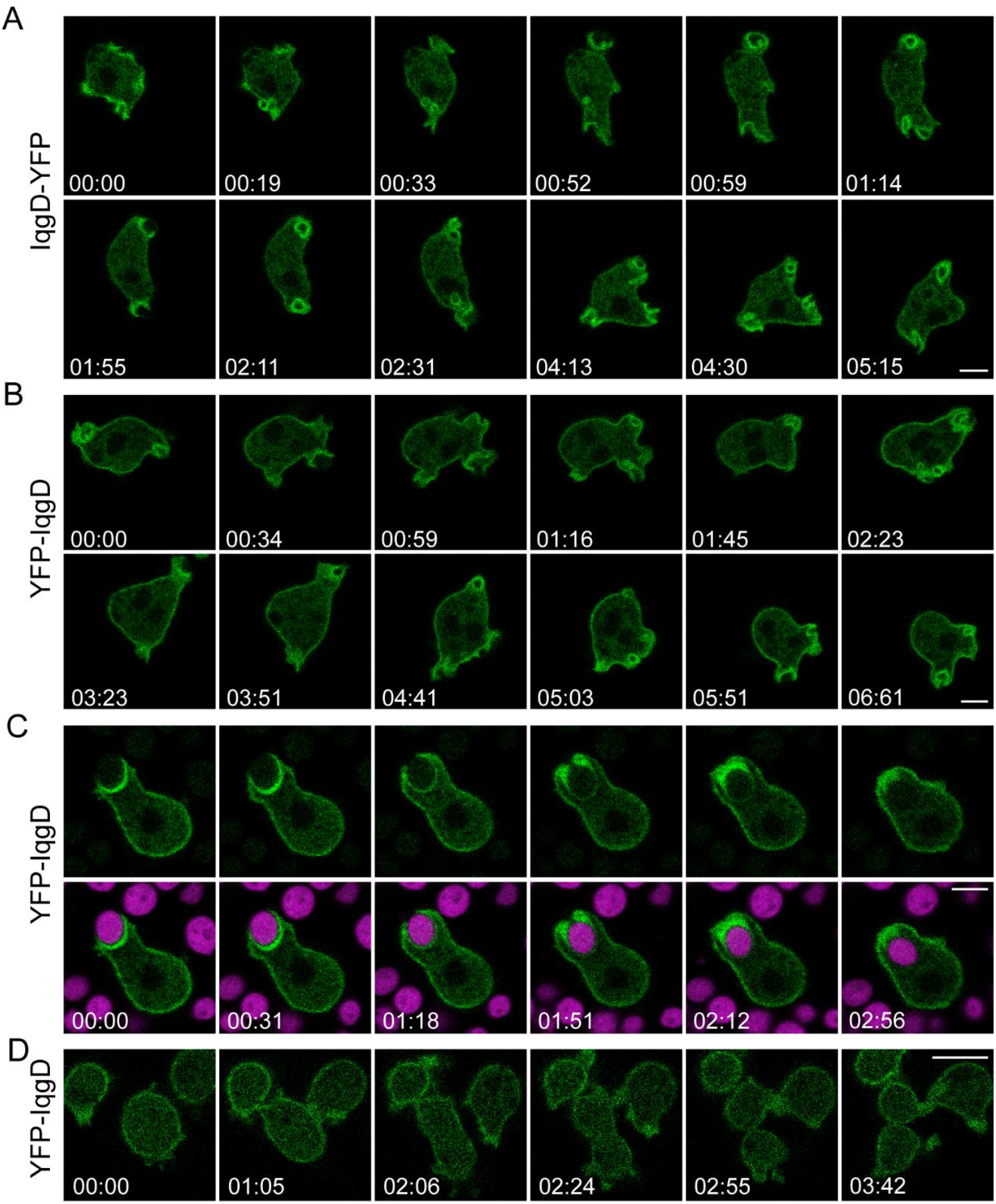
IqgD localizes over the entire cell cortex with strong enrichment at macropinosomes and phagosomes. Image sequences of growth-phase AX2 cells extrachromosomally expressing IqgD-YFP (A) and YFP-IqgD (B) during random movement and macropinocytosis, YFP-IqgD during phagocytosis of TRITC-labeled yeast (magenta) (C), and cytokinesis (D). Time is given in min:sec format. Scale bars: 5 μm (A-C); 10 μm (D). A-D correspond to Movies S1-S4.

### IqgD interacts with F-actin via its CHD

To investigate whether IqgD interacts with F-actin, we performed an actin co-sedimentation assay using bacterially purified protein. Because full-length IqgD was poorly expressed in bacterial cells, we purified an N-terminal fragment (aa 1-575) containing the CHD. The actin co-sedimentation assay using this fragment showed no detectable interaction with F-actin (Fig. S2A). Next, we performed co-immunoprecipitation with an anti-IqgD antibody raised against the same N-terminal fragment to test whether actin binds to endogenous IqgD. However, actin bound substantially to the protein A-Sepharose beads even in the absence of the anti-IqgD antibody, making this method unsuitable for detecting IqgD-actin interaction (Fig. S2B). Therefore, we used indirect approaches to assess whether IqgD associates with F-actin. First, we monitored the localization of YFP-IqgD during disruption of the actin cytoskeleton with Latrunculin A (Fig. 2A). Shortly after Latrunculin A addition, the cells rounded up, and within five minutes YFP-IqgD largely detached from the cortex. After 20 minutes, the signal became completely cytosolic. After washing out the drug, YFP-IqgD reappeared at the cell cortex. We then examined the localization of an IqgD variant lacking the CHD, IqgD_ΔCHD (Fig. 2B). This truncated protein was diffusely cytosolic and failed to accumulate in macropinocytic cups. These experiments showed that IqgD localization to F-actin-rich structures requires an intact cortical actin network and depends on its CHD. Prompted by these results, we repeated the actin co-sedimentation assay with an N-terminal fragment (aa 1-592) of IqgD produced in insect cells. Using this protein produced in eukaryotic cells, we detected direct binding to F-actin in the co-sedimentation assay (Fig. 2C). Additionally, we compared F-actin distribution in cells with different IqgD levels. Immunofluorescence with an anti-actin antibody showed similar actin distribution in wild-type, IqgD-deficient, and IqgD-overexpressing cells (Fig. S2C). Quantification of the F- to G-actin ratio showed no difference between wild-type and IqgD-deficient cells (Fig. S2D). However, the F-actin level was significantly increased in IqgD-overexpressing cells.

**Figure 2.**
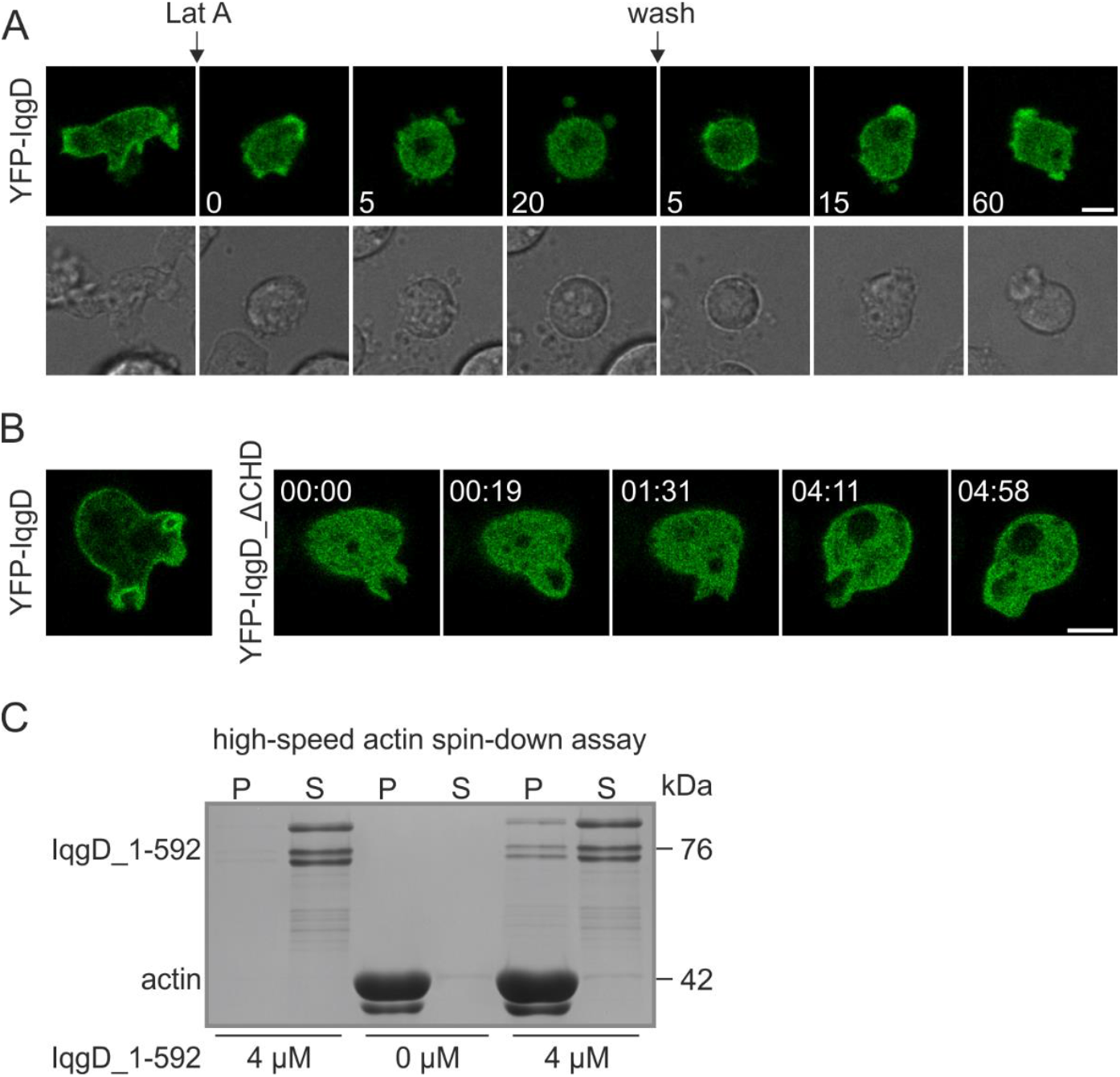
Cortical localization of IqgD requires interaction of the CHD with an intact cortical F-actin network. (A) Wild-type cells expressing YFP-IqgD before, during, and after treatment with 5 *µ*M Latrunculin A. The numbers to the right of the *Lat A* label indicate minutes after addition of latrunculin. The numbers to the right of the *wash* label indicate minutes after washing the cells with fresh medium. Scale bar: 5 *µ*m. (B) Image sequence of a representative vegetative IqgD-deficient cell expressing YFP-IqgD_ΔCHD. Time is given in min:sec format. Scale bar: 5 µm. The sequence corresponds to Movie S7. (C) The Coomassie Brilliant Blue (CBB)-stained PAG of the high-speed actin spin-down assay shows that IqgD_1-592 produced in insect cells co-sediments with F-actin. *P* and *S* indicate the pellet and supernatant, respectively.

### IqgD interacts with active Rac1 GTPases in living cells

The localization of IqgD to macropinosomes and phagosomes suggests its involvement in macroendocytosis. Additionally, IqgD harbors a GRD, which mediates interactions between IQGAPs and Ras superfamily GTPases (30). Therefore, we tested the interaction of IqgD with Ras and Rho family GTPases known to be involved in macroendocytosis. A yeast two-hybrid assay with full-length IqgD showed weak interactions with the GTPases Rac1A and Rac1C (Fig. S3A), whereas the assay using only the GRD of IqgD detected interactions with the constitutively active forms of all three Rac1 GTPases (Fig. 3A and Fig. S3B). After establishing that IqgD largely colocalizes with the probe specific for active Rac1 GTPases (43) (Fig. 3B), we performed a GST-Rac1A pull-down assay and found that IqgD binds to both GTPγS- and GDP-loaded Rac1A, binding even somewhat more strongly to the GDP-loaded GTPase (Fig. 3C). This was corroborated in an independent GST-Rac1A pull-down using constitutively active and constitutively inactive Rac1A mutants (Fig. S4). To evaluate direct interactions between IqgD and Rac1 GTPases in live *D. discoideum* cells, we used the bimolecular fluorescence complementation (BiFC) assay. *IqgD* knockout (*iqgD*^*L1-*^, see below) cells were transfected with an expression vector allowing simultaneous expression of IqgD fused at its N-terminus to the C-terminal part of the fluorescent protein Venus (VC-IqgD) and the respective Rac1 GTPase fused at its N-terminus to the N-terminal part of Venus (VN-Rac1). Complementation of the Venus protein due to interaction between IqgD and the tested GTPase resulted in a strong fluorescent signal for the wild-type forms of all three Rac1 GTPases (Fig. 3D). To assess the relevance of the nucleotide status of the interacting GTPase, we co-expressed VC-IqgD with VN-Rac1A/1C GTPases as mutant variants: two constitutively active (G12V and Q61L) and one constitutively inactive (T17N) variant. A fluorescent signal was detected only with the constitutively active Rac1 variants. However, the signal intensity was markedly weaker compared to the wild-type GTPases and was observed in significantly fewer cells (Fig. 3E). The strong cortical BiFC signal obtained with wild-type GTPases indicated that GTPase cycling is required and suggested that interactions between IqgD and Rac1 GTPases occur throughout the cell cortex, consistent with the localization of YFP-IqgD. However, closer inspection revealed subtle differences in the distribution of YFP-IqgD and the BiFC signals. Specifically, YFP-IqgD was enriched in the macropinocytic and phagocytic cups, whereas the Venus signals, particularly those from the interaction of wild-type Rac1A and Rac1B with IqgD were stronger in the posterior regions of the cell cortex and weaker at the macropinocytic cups (compare Fig. 1B and 1C with *the upper and middle panels* in Fig. 3D). In contrast, the Venus signal resulting from the interaction of Rac1C with IqgD appeared equally strong at the macropinosomes and in the posterior regions of the cortex (Fig. 3D, *lower panel*). To verify this observation, we expressed all three VN-VC protein pairs together with the probe for active Ras as a marker for macropinocytic cups and measured the Venus signal intensity at the cup base and in the retractive region of the cell cortex (Fig. S5 A-C). The BiFC signals reflecting interactions between IqgD and Rac1A or Rac1B were about 80% stronger at the posterior parts of the cell cortex than at the base of the macropinocytic cups (Fig. S5D). By contrast, the interaction between IqgD and Rac1C showed no preference for any particular cortical region, based on the corresponding BiFC signal.

**Figure 3.**
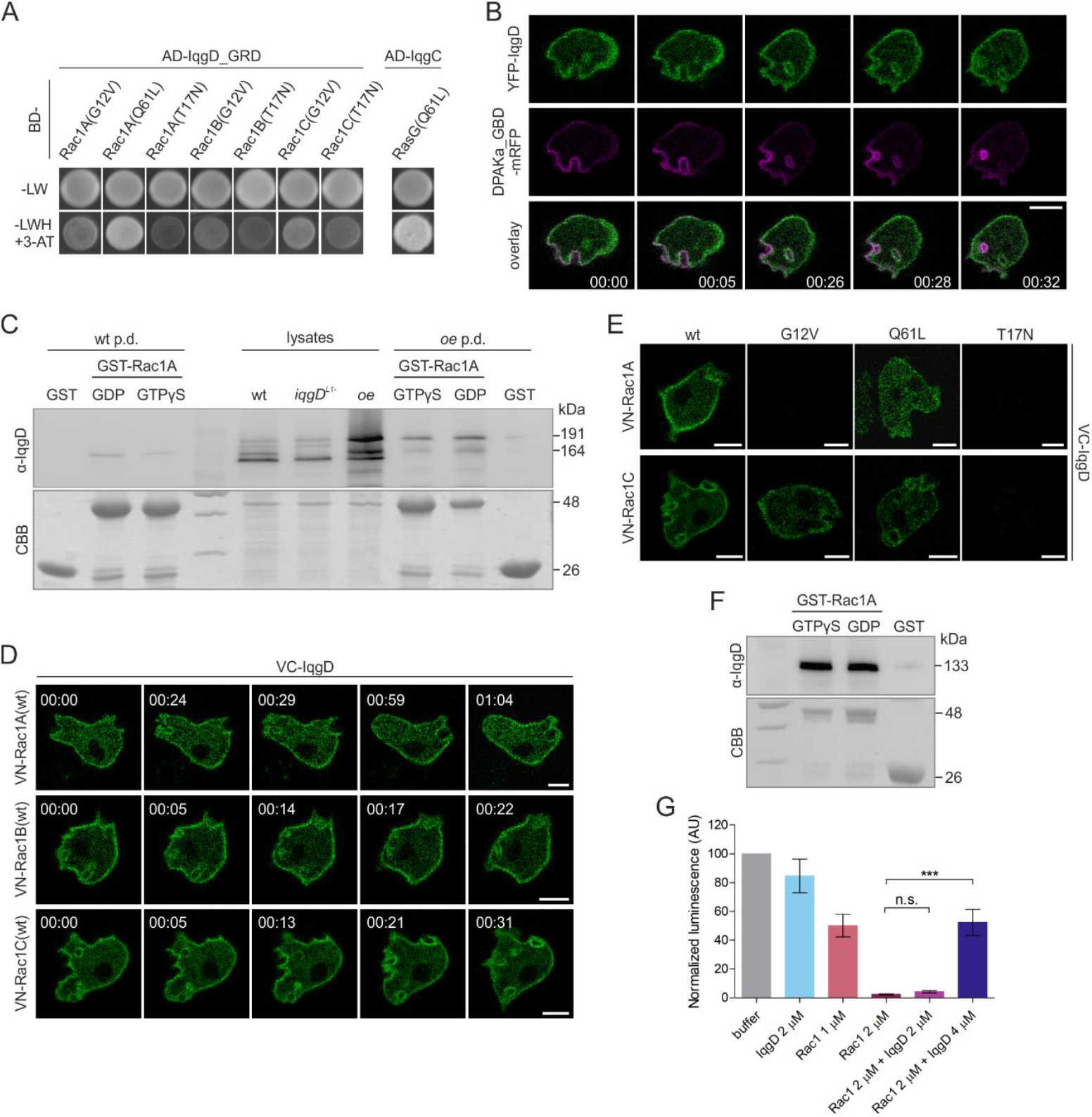
IqgD interacts with active Rac1 GTPases in living cells and protects their active state. (A) The GRD of IqgD interacts with active forms of Rac1 GTPases in the yeast two-hybrid assay. For details, see Fig. S3. (B) Image sequence of a wild-type vegetative cell expressing YFP-IqgD and the probe for active Rac1 GTPases, DPAKa_GBD-mRFP. (C) Anti-IqgD blot of GST-Rac1A pull-down with lysates of wild-type cells (*wt p*.*d*.) and wild-type cells expressing YFP-IqgD (*oe p*.*d*.) shows that both endogenous IqgD (164 kDa) and exogenous YFP-IqgD (191 kDa) interact with GTPγS- and GDP-loaded Rac1A. The GST pull-down was performed as a negative control. Lysate from *iqgD*^*L1*-^ cells was loaded in the center of the membrane to facilitate recognition of a specific band. See also Fig. S6A. (D) Image sequences of vegetative *iqgD*^*L1*-^ cells expressing VC-IqgD with VN-Rac1A(wt) (*upper panel*), VN-Rac1B(wt) (*middle panel*) or VN-Rac1C(wt) (*lower panel*). (E) Images of *iqgD*^*L1*-^ cells expressing VC-IqgD with VN-Rac1A(wt/G12V/Q61L/T17N) (*upper panel*) or VN-Rac1C(wt/G12V/Q61L/T17N) (*lower panel*). (F) Anti-IqgD blot of GST-Rac1A pull-down with IqgD_311-1385 purified from insect cells confirms direct binding of IqgD to Rac1A. The GST pull-down was performed as a negative control. (G) The results of the GAP assay show that IqgD_311-1385 added in double concentration to human Rac1 reduces its intrinsic GTPase activity (compare *Rac1 2 *µ*M* with *Rac1 2 *µ*M + IqgD 4 *µ*M*). An equimolar concentration of IqgD_311-1385 was not sufficient to significantly increase luminescence, i.e. to reduce GTP hydrolysis by Rac1. Data are from 3 independent experiments (mean ± SD). In (B) and (D), time is given in min:sec format. Scale bars in (B), (D), and (E): 5 μm. (B) and (D) correspond to Movies S8 and S9-11, respectively. In (C) and (F), CBB-stained PAGs are displayed underneath the blots. Statistical analysis in (G) was performed using one-way ANOVA followed by Tukey’s multiple comparison test. The significance level was set at 5%. n.s. – not significant, *** P < 0.001.

Finally, after establishing that IqgD interacts with active Rac1 GTPases in living cells, we examined whether it acts as a GAP for Rac1. To this end, we attempted to purify several truncated IqgD variants containing the GRD with or without the RGCt domain from *E. coli*, but were unable to obtain a soluble protein. Therefore, we again produced the protein from insect cells and successfully obtained IqgD_311-1385, the truncated variant containing both the GRD and RGCt domains. To confirm that the protein is biologically functional, we repeated the GST-Rac1A pull-down assay with IqgD_311-1385 and showed that it binds strongly to both GTPγS- and GDP-bound forms of Rac1A with similar affinities (Fig. 3F). We then measured the GTPase activity of human Rac1 in the presence and absence of IqgD_311-1385 and showed that IqgD has no GAP activity toward Rac1 (Fig. 3G). Moreover, similar to its mammalian counterparts, IqgD inhibited the intrinsic GTPase activity of Rac1, thus stabilizing its active state.

### IqgD binding to cortexillins and Rac1 is mutually exclusive

A recent study showed that the GRD, RGCt, and CT domains of human IQGAP1 are involved in binding Cdc42 and Rac1 (21). Therefore, we investigated the importance of the GRD and RGCt domains of IqgD in binding *D. discoideum* Rac1 GTPases. We chose Rac1C for this analysis because our BiFC results indicated that Rac1C interacted equally strongly with IqgD at both the anterior and posterior cortical regions (Fig. S5C and D). We transfected *iqgD*^*L1*-^ cells (see below) with three additional BiFC constructs to express VC fused with either IqgD lacking the GRD, IqgD lacking the RGCt, or the GRD alone, together with VN-Rac1C(wt). In contrast to cells transfected with the BiFC construct encoding full-length IqgD (VC-IqgD_FL), which displayed cortical fluorescence in most cells, cells expressing IqgD without the GRD (VC-IqgD_ΔGRD) or the GRD alone (VC-IqgD_GRD) showed only weak Venus fluorescence in a small fraction of cells (Fig. 4A). No fluorescence was observed in cells expressing IqgD without the RGCt domain (VC-IqgD_ΔRGCt). These results suggest that both the RGCt and the GRD contribute to Rac1 binding. Next, we examined the localization of truncated IqgD variants lacking either the GRD or the RGCt domain. Interestingly, deletion of either domain resulted in a clear separation of IqgD localization between front and back structures. These findings strongly suggest that the RGCt domain is required for recruiting IqgD to macropinocytic cups, whereas the GRD is important for its localization to retracting posterior structures (Fig. 4B). We previously showed that Rac1 GTPases play a dual role in regulating the actin cytoskeleton in *D. discoideum* (38). At the cell front, Rac1-GTP promotes dendritic actin nucleation by activating WASP family proteins upstream of the Arp2/3 complex (34). At the rear, Rac1-GTP promotes the formation of a tetrameric complex consisting of DGAP1 or GAPA and a dimer of the actin-binding proteins, the cortexillins, which bundle actin filaments (35, 44). The localization of IqgD_ΔGRD closely resembles that of a DPAKa_GBD probe specific for active Rac1 GTPases at the front, whereas IqgD_ΔRGCt adopts the same posterior localization as the Rac1 interactor DGAP1 and cortexillin I (38). Therefore, we investigated whether IqgD, similar to DGAP1 and GAPA, also interacts with the cortexillins. In a co-immunoprecipitation assay using an anti-IqgD antibody, both cortexillin I (CI) and II (CII) co-precipitated with endogenous IqgD (Fig. 4C). To test whether the binding is direct, we again performed a BiFC assay. *IqgD*^*L1*-^ cells were transfected with VN-CI or VN-CII together with either VC-IqgD_FL, VC-IqgD_ΔGRD, or VC-IqgD_ΔRGCt. We observed a strong fluorescent signal in the posterior cortical regions of cells expressing either cortexillin with VC-IqgD_FL (Fig. 4D). However, in cells expressing truncated IqgD variants, no fluorescence was detected, suggesting that both the GRD and RGCt domains are required for binding to the cortexillins. Previously, it was shown that binding of DGAP1 or GAPA to the cortexillin dimer requires the association of active Rac1A with the IQGAP-related protein (35). Since IqgD binds directly to Rac1A (Fig. 3D and F) as well as to cortexillin I and II (Fig. 4D), we wondered whether IqgD might also participate in the formation of a tetrameric complex with Rac1 and the cortexillin dimer. To test this, we bound IqgD_311-1385 to GST-Rac1A as shown in Fig. 3F and added lysate of *dgap1*^*-*^*/gapA*^-^ cells to such immobilized IqgD. However, we did not detect CI or CII in the complex with Rac1-bound IqgD (Fig. 4E). This is consistent with previous findings showing that cortexillins cannot be detected in a GST-Rac1A pull-down using lysates from *dgap1*^*-*^*/gapA*^-^ cells (45). To confirm that truncated IqgD_311-1385 can interact with the cortexillins, we performed a BiFC assay. Fluorescence was detected in cells co-expressing VC-IqgD_311-1385 and VN-CI or VN-CII (Fig. 4D), although the signal was weaker compared to full-length IqgD, probably because IqgD associates less efficiently with cortexillins in the absence of its CHD domain.

**Figure 4.**
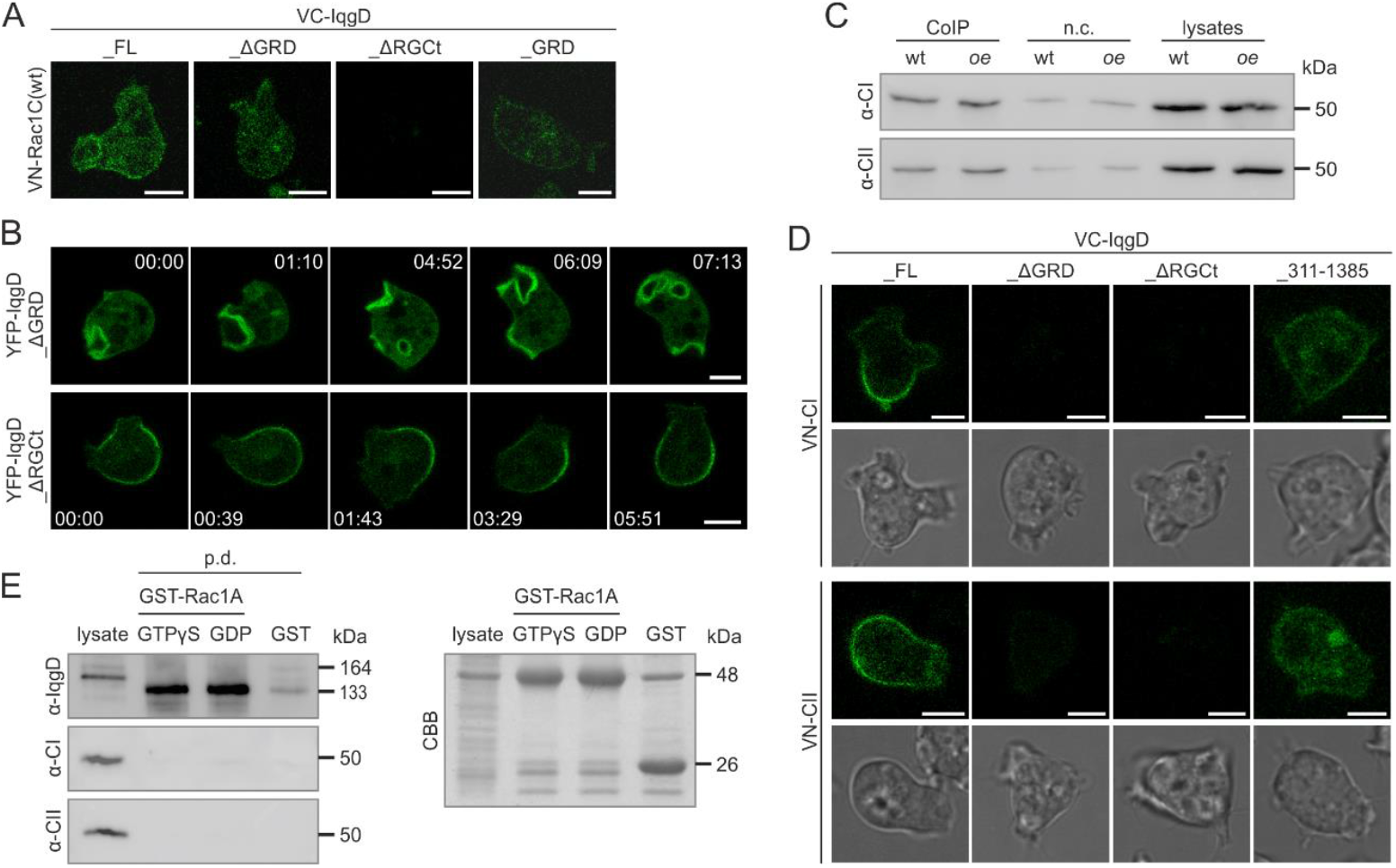
The GRD and RGCt domains of IqgD are required for binding to Rac1 and cortexillin. (A) Images of vegetative *iqgD*^*L1*-^ cells expressing VN-Rac1C(wt) with VC-IqgD_FL (fulllength), VC-IqgD_ΔGRD, VC-IqgD_ΔRGCt, or VC-IqgD_GRD. (B) Image sequences of vegetative *iqgD*^*L1*-^ cells expressing YFP-IqgD_ΔGRD (*upper panel*) and YFP-IqgD_ΔRGCt (*lower panel*). (C) Anti-CI (*upper membrane*) and anti-CII (*lower membrane*) blots of co-immunoprecipitation of cortexillins I and II with IqgD from lysates of wild-type (wt) and IqgD-overexpressing (*oe*) cells (*CoIP*). Co-immunoprecipitation with non-specific IgG was performed as a negative control (*n*.*c*.). (D) Images of *iqgD*^*L1*-^ cells co-expressing either VN-CI (*two upper panels*) or VN-CII (*two lower panels*) with VC-IqgD_FL, VC-IqgD_ΔGRD, VC-IqgD_ΔRGCt, or VC-IqgD_311-1385. (E) Anti-IqgD blot (*upper membrane*) of GST-Rac1A pull-down shows that IqgD_311-1385 bound to both GTPγS- and GDP-loaded Rac1A. However, CI or CII from lysates of *dgap1*^*-*^*/gapA*^-^ cells did not associate with IqgD_311-1385, as shown by the anti-CI (*middle membrane*) and anti-CII (*lower membrane*) blots (*left*). The CBB-stained PAG of GST proteins used in the pull-down assay is shown (*right*). In (B), time is given in min:sec format. Scale bars in (A), (B) and (D): 5 μm. (B) corresponds to Movies S12 and S13.

### The axenic strain AX2-214 harbors two copies of the *iqgD* gene

To investigate the cellular function of IqgD, we aimed to generate IqgD-deficient cells by inactivating *iqgD* using homologous recombination. Although a unique recombination event at the target locus (*iqgD*^*L1*^) was confirmed by PCR, Southern blotting, and immunoblotting with a polyclonal anti-IqgD antibody raised against the N-terminal fragment of IqgD (Fig. S6A), we discovered a second copy of the gene (designated *iqgD*^*L2*^) in the *D. discoideum* genome (Fig. S6B). However, this second locus appears to be poorly expressed, as evidenced by strikingly reduced transcript levels in *iqgD*^*L1*-^ cells (Fig. S6C). To determine whether the presence of an alternative *iqgD* locus is intrinsic to strain AX2-214 or a consequence of cell propagation in our laboratory, we obtained a vial of AX2-214 spores stored in 1992. We repeated the homologous recombination and confirmed the inactivation of *iqgD*^*L1*^ by PCR (Fig. S6D) and western blot (Fig. S6A). However, PCR with primers specific for *iqgD*^*L2*^ confirmed the presence of the second locus (Fig. S6E). Sequencing the first 2.47 kb of *iqgD*^*L2*^ using genomic DNA from cells with inactivated *iqgD*^*L1*^ (*iqgD*^*L1-*^) revealed that the sequence is identical to that of *iqgD*^*L1*^ except that all introns are absent. Therefore, *iqgD*^*L2*^ appears to be a weakly active retrogene (Fig. S6F). To formally exclude residual expression from the second locus, we excised the bsr resistance cassette from *iqgD*^*L1*-^ cells by transient expression of Cre recombinase and subsequently inactivated *iqgD*^*L2*^ by homologous recombination. Successful inactivation of *iqgD*^*L2*^ was confirmed by PCR and sequencing (Fig. S6G).

### IqgD is required for normal growth on bacterial lawns

Since IqgD is distributed throughout the cortex with strong enrichment in macroendocytic cups, we compared the efficiency of fluid and particle uptake, as well as other actin-driven processes, in wild-type and IqgD-deficient cells. As *iqgD*^*L1*-^ cells express negligible amounts of IqgD (Fig. S6C), we used them for most experiments. First, we examined the growth of *iqgD*^*L1*-^ cells in suspension. Although they grew slightly faster than wild-type cells, the difference was not statistically significant (Fig. 5A). Consistently, *iqgD*^*L1*-^ cells did not show increased macropinocytosis efficiency (Fig. 5B), and the size of their macropinosomes was not significantly different from that of wild-type cells (Fig. 5C). Growth in suspension and macropinosome size were compared for three and two *iqgD*^*L1*-^ clones, respectively, to demonstrate the consistency of the phenotype among independent clones. Next, we analyzed the growth of *iqgD*^*L1*-^ cells on a lawn of *K. aerogenes* and found that they formed markedly smaller plaques compared to wild-type cells (Fig. 5D). Assuming that IqgD functions as a Rac1 effector, we compared the growth of *iqgD*^*L1*-^ and *rac1A*^-^ cells. We used *rac1A*^-^ cells because Rac1A is the most highly expressed Rac1 isoform in vegetative AX2 cells (46, 47). The *rac1A*^-^ cells were generated by homologous recombination, and successful inactivation of the *rac1A* gene was confirmed by PCR and Southern blot (Fig. S7). As expected, *rac1A*^-^ cells grew significantly slower than both wild-type and *iqgD*^*L1*-^ cells on bacterial lawns (Fig. 5D). Because slow growth on bacterial lawns is most likely caused by inefficient phagocytosis of bacteria, we examined the uptake of *E. coli* strain DH5α in suspension. Unexpectedly, however, neither *iqgD*^*L1*-^ nor *rac1A*^-^ cells showed reduced uptake of bacteria (Fig. 5E). To exclude potential effects arising from the use of different bacterial species for growth on bacterial lawns and phagocytosis in suspension, we repeated both assays with *E. coli* strain B/2, which is considered suitable for phagocytosis under both conditions. However, once again we observed significantly reduced growth rates for *iqgD*^*L1*-^ and *rac1A*^-^ cells compared to wild-type cells on bacterial lawns (Fig. 5F), whereas *iqgD*^*L1-*^, *rac1A*^*-*^, IqgD-overexpressing, and wild-type cells all showed similar phagocytosis efficiency in suspension (Fig. 5G). We also included *iqgD*^*L1-/L2*-^ double knockout cells and found no difference in plaque size between *iqgD*^*L1*-^ and *iqgD*^*L1-/L2*-^ cells (Fig. 5F).

**Figure 5.**
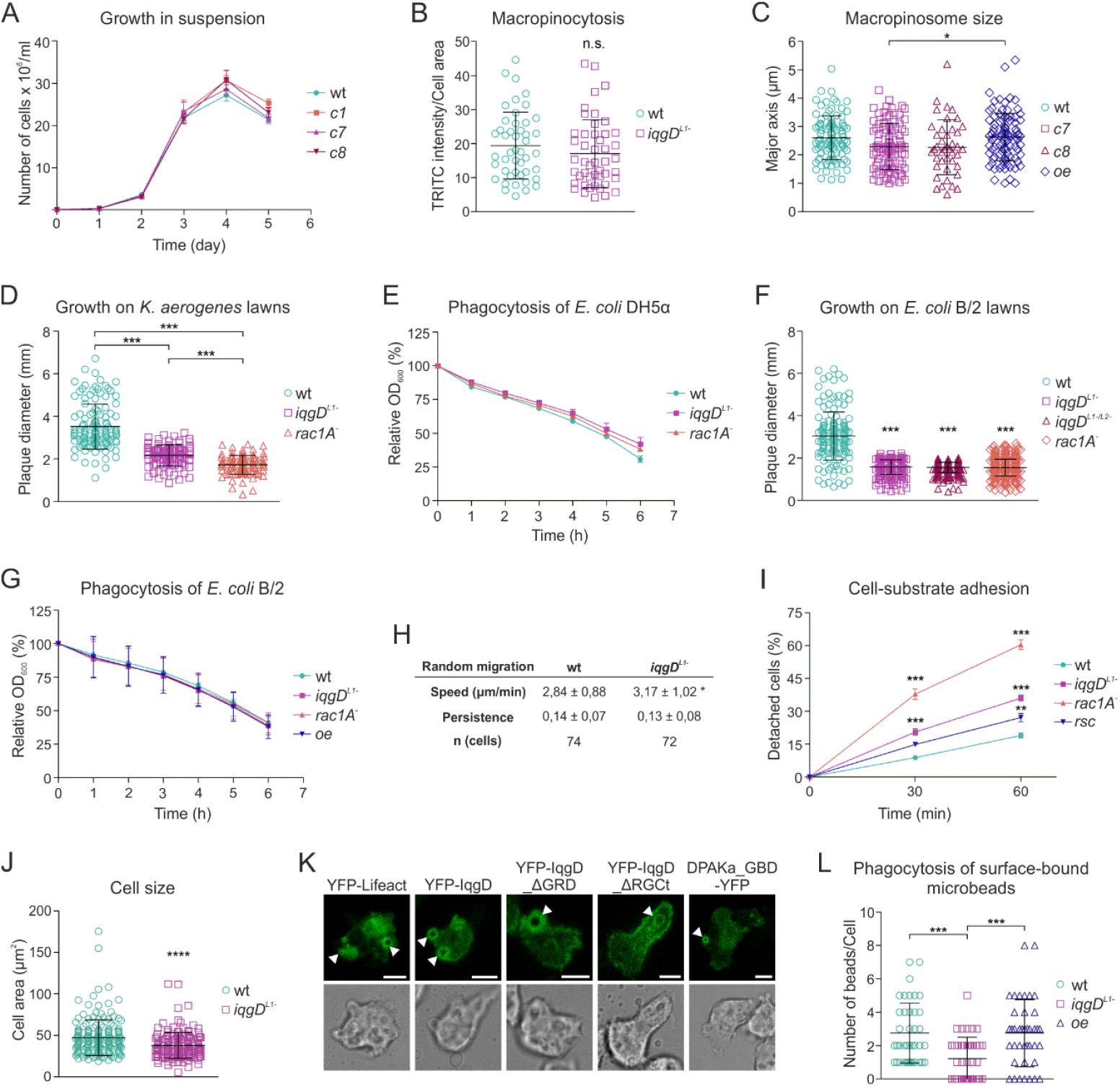
*IqgD*^*L1*-^ cells exhibit a severe growth defect on bacterial lawns due to impaired phagocytosis of surface-bound particles and poor adhesion to the substrate. (A) *IqgD*^*L1*-^ cells (clones *c1, c7*, and *c8*) grow at a rate similar to wild-type (wt) cells in shaken liquid culture medium (n ≥ 6, mean ± SEM). (B) There is no statistically significant difference in fluid uptake between wt and i*qgD*^*L1*-^ cells. Data show the results of one of four experiments in which the ratios of TRITC intensity to cell area were: 19.4 ± 9.9 for wt, n = 46; and 17 ± 10 for *iqgD*^*L1-*^, n = 47 (mean ± SD). (C) Macropinosomes of *iqgD*^*L1*-^ cells (*c7* and *c8*) are slightly smaller than those of wt cells, while cells expressing YFP-IqgD (*oe*) have macropinosomes indistinguishable from wt cells. The length of the macropinosome major axis determined from at least 3 independent experiments was: 2.6 ± 0.8 *µ*m for wt, n = 97; 2.3 ± 0.8 *µ*m for *iqgD*^*L1-*^, *c7*, n = 92; 2.3 ± 1 *µ*m for *iqgD*^*L1-*^, *c8*, n = 42; and 2.6 ± 0.8 for *oe*, n = 95 (mean ± SD). (D) *IqgD*^*L1*-^ and *rac1A*^-^ cells grow significantly slower than wt cells on solid plates with *K. aerogenes* as the food source. The diameters of plaques from 4 independent experiments measured on day 4 were: 3.5 ± 1.1 mm for wt, n = 123; 2.2 ± 0.5 mm for *iqgD*^*L1-*^, n = 98; and 1.7 ± 0.4 mm for *rac1A*^*-*^, n = 93 (mean ± SD). (E) *IqgD*^*L1*-^ and *rac1A*^-^ cells show similar efficiency in phagocytosis of *E. coli* DH5α from shaken suspension as wt cells (n ≥ 3, mean ± SEM). (F) *IqgD*^*L1-*^, *iqgD*^*L1-/L2-*^, and *rac1A*^-^ cells grow significantly more slowly than wt cells on solid plates with *E. coli* B/2 as the food source. The diameters of plaques from 4 independent experiments measured on day 5 were: 3 ± 1.1 mm for wt, n = 154; 1.6 ± 0.4 mm for *iqgD*^*L1-*^, n = 231; 1.6 ± 0.2 mm for *iqgD*^*L1-/L2-*^, n = 356; and 1.5 ± 0.4 mm for *rac1A*^*-*^, n = 438 (mean ± SD). (G) *IqgD*^*L1-*^, *rac1A*^*-*^, and *oe* cells show similar efficiency in phagocytosis of *E. coli* strain B/2 from shaken suspension as wt cells (n ≥ 5, mean ± SEM). (H) The speed of randomly moving *iqgD*^*L1*-^ cells is slightly increased compared to wt cells, while persistence is not altered. Data are from 4 independent experiments (mean ± SD). (I) *IqgD*^*L1*-^ and *rac1A*^-^ cells are less strongly attached to the substrate compared to wt cells in a detachment assay (n ≥ 4, mean ± SEM). Expression of exogenous YFP-IqgD in *iqgD*^*L1*-^ cells (*rsc*) partially rescues their adhesion defect. (J) *IqgD*^*L1*-^ cells harvested from the vegetative zone of the plaque on *K. aerogenes* are significantly smaller than their wild-type counterparts. The cell areas from 3 independent experiments were: 47.2 ± 21.4 μm^2^ for wt, n = 174; and 37.8 ± 15.5 μm^2^ for *iqgD*^*L1-*^, n = 154 (mean ± SD). (K) Confocal images of the ventral surface of wild-type cells expressing YFP-Lifeact, YFP-IqgD, YFP-IqgD_ΔGRD, YFP-IqgD_ΔRGCt, and DPAKa_GBD-YFP. The cells were attached to the glass surface covered with silicone grease droplets of various sizes in the micrometer range. Scale bars: 5 μm. Images of cells expressing YFP-Lifeact, YFP-IqgD and DPAKa_GBD-YFP correspond to Movies S15-S17. (L) *IqgD*^*L1*-^ cells are less efficient than wild-type cells in phagocytosis of surface-bound 1 μm beads. The numbers of ingested beads in 30 minutes were: 2.8 ± 1.8 for wt, n = 40; 1.2 ± 1.3 for *iqgD*^*L1-*^, n = 40; and 2.8 ± 2 for *oe* cells, n = 40 (mean ± SD). Statistical analyses were performed using: two-way RM ANOVA with Bonferroni posttests in (A), (E), (G), and (I); Mann-Whitney test in (B), (H), and (J); Kruskal-Wallis test followed by Dunn’s multiple comparison test for pairwise comparisons in (C), (D), (F), and (L). The significance level was set at 5% for all analyses. * P < 0.05, ** P < 0.01, *** P < 0.001, **** P < 0.0001.

### IqgD is required for phagocytosis of surface-attached particles

As *iqgD*^*L1*-^ cells showed no cytokinesis defect when grown on Petri dishes, we examined cell motility and adhesion to identify other possible reasons for their poor growth on bacterial lawns. Randomly moving vegetative *iqgD*^*L1*-^ cells displayed only a slightly increased speed and unchanged directional persistence compared to wild-type cells (Fig. 5H). However, their adhesion to the substrate was significantly reduced compared to wild-type cells (Fig. 5I). *Rac1A*^-^ cells exhibited an even more pronounced defect in cell-substrate adhesion. Using TIRF microscopy, we then investigated whether IqgD localizes to adhesion foci, but instead found it to be evenly distributed throughout the ventral cortex (Movie S14). Impaired adhesion during growth on bacterial lawns could affect cytokinesis, phagocytosis, or motility, even if these processes appear unaffected under other conditions. Therefore, we harvested cells from the vegetative zone of the spreading plaque, fixed them, and stained them with DAPI. Most wild-type and *iqgD*^*L1*-^ cells were mononucleated, with a very small proportion of binucleated cells that did not differ statistically between the cell lines (wt 2.3%, n = 174; *iqgD*^*L1*-^ 0.65%, n = 154). However, the measured cell area of *iqgD*^*L1*-^ cells was significantly smaller than that of wild-type cells (Fig. 5J). Based on these results, we hypothesized that *iqgD*^*L1*-^ cells have a specific defect in phagocytosis of bacteria attached to the agar surface, possibly involving a mechanism similar to that used by macrophages to detach and internalize surface-bound particles (48). Macrophages form an F-actin-rich circular adhesive structure, termed the phagocytic adhesion ring (PAR), around the particle at the cell-substrate interface, which provides the force for its detachment from the surface. To investigate whether a similar structure exists in *Dictyostelium*, we plated wild-type cells expressing an F-actin probe on a glass surface covered with silicone grease droplets. Interestingly, F-actin encircled the droplets underneath the cell in a manner similar to PARs in macrophages (Fig. 5K and Movie S15). We then found that IqgD and active Rac1 also localize to this structure (Fig. 5K, and Movies S16 and S17). Notably, both IqgD_ΔGRD and IqgD_ΔRGCt were able to localize to these ventral rings, although the localization of IqgD lacking the RGCt was observed less frequently and generally with a weaker signal (Fig. 5K).

To investigate the relevance of IqgD in the phagocytosis of surface-bound particles, we adapted a recently described protocol developed for macrophages (48). Wild-type, *iqgD*^*L1-*^, and IqgD-overexpressing cells were plated on a poly-L-lysine-coated glass surface to which 1 μm fluorescent beads had been immobilized. After 30 minutes, the cells were fixed, and the number of ingested particles was determined using confocal microscopy. Notably, *iqgD*^*L1*-^ cells phagocytosed significantly fewer particles than wild-type cells (Fig. 5L), strongly suggesting that their reduced growth on bacterial lawns is caused by inefficient phagocytosis of bacteria attached to the surface. Therefore, we identified IqgD as a Rac1 effector that interacts with F-actin and is important for efficient growth of *D. discoideum* on bacterial lawns by facilitating the uptake of bacteria attached to the surface of the agar plate.

## Discussion

In this study, we set out to define the cellular role of the IQGAP-related protein IqgD from *D. discoideum*. Given that IqgD contains CHD, GRD, and RGCt domains that, in human IQGAP1, mediate interactions with F-actin and the Rho GTPases Cdc42 and Rac1, we investigated whether IqgD similarly interacts with F-actin and Rho-family GTPases in *D. discoideum*. We show that IqgD localizes to F-actin-rich structures such as the cell cortex and macroendocytic cups, and that its cortical localization depends on an intact actin cortex and the presence of the CHD. Consistently, we found that IqgD binds directly to F-actin, although this interaction appears to be of relatively low affinity, at least when using rabbit skeletal muscle actin, as assessed by co-sedimentation assays using an N-terminal fragment containing the CHD. In contrast, Hoeprich et al. reported that a monomeric N-terminal fragment of human IQGAP1 containing the CHD is sufficient for stable binding along filament sides (26). Its dwell time appears to be comparable to that of the dimeric full-length protein, and both full-length human IQGAP1 and the N-terminal fragment bind to filament sides with high affinity. By contrast, the relatively low binding affinity of IqgD may be explained by the presence of a different type of CHD compared with that in IQGAP1. Mammalian IQGAPs contain a single type 3 CHD, whereas IqgD harbors two fimbrin-type CHDs, CHf1 and CHf2 (49, 50). Fimbrins are potent actin-bundling proteins that contain two CHD duplexes, which together constitute two actin-binding domains (ABDs) (49). In general, both fimbrins and IQGAPs exhibit moderate to high binding affinity for F-actin sides (26, 51–55). However, in contrast to canonical fimbrins, IqgD harbors only one duplex and was therefore originally referred to as a fimbrin-related RasGAP (50). Of note, it is well established that the ABDs of fimbrin differ in their binding affinities for F-actin, with the isolated ABD2 binding substantially more strongly than ABD1 (55–57). Accordingly, the presence of only the first two CHDs of fimbrin, corresponding to ABD1, in IqgD may account for its relatively low affinity for F-actin. Unfortunately, the low yield of IqgD_1-592 from insect cells precluded assessment of its bundling activity in a low-speed actin spin-down assay.

We further show that IqgD interacts with the *D. discoideum* orthologs of human Rac1, namely the GTPases Rac1A, Rac1B, and Rac1C (58). Although IqgD bound more strongly to inactive Rac1A in GST-Rac1A pull-down assays, BiFC analysis showed that it interacts exclusively with active Rac1 GTPases in living cells. This, along with the finding that IqgD does not function as a GAP for Rac1 GTPases, suggests that it acts as a Rac1 effector. Consistent with this notion, the phenotype of *iqgD*^*L1*-^ cells overlaps with that of *rac1A*^-^ cells. However, cell adhesion and growth on *K. aerogenes* lawns are more strongly impaired in the absence of Rac1A than in the absence of IqgD, probably because Rac1A functions as an upstream regulator that signals through multiple downstream effectors in distinct pathways.

IqgD is present in diverse F-actin-based structures, including anterior protrusive structures such as macropinocytic cups, where it is highly enriched, as well as posterior retractive structures. The CHD is essential for overall cortical localization through its ability to bind F-actin, as IqgD variants lacking the CHD are largely cytosolic. However, truncated IqgD variants that retain the CHD but lack either the GRD or the RGCt domain are restricted to anterior or posterior cortical regions, respectively. Thus, while the CHD mediates general cortical attachment, the GRD and RGCt domains further specify distinct cortical localizations. The RGCt domains of human IQGAP1 and IQGAP2 exhibit approximately 100-fold higher affinity for active than for inactive Cdc42, whereas their GRDs bind Cdc42 with low affinity independent of the nucleotide state (2). Based on these findings, the RGCt domain has been proposed to function as the actual effector or GTPase-binding domain (GBD) of IQGAP, with the GRD serving mainly as a scaffold for the GTPase (2, 21). Consistent with this model, we show that both the GRD and RGCt domains of IqgD contribute to Rac1 binding, with the RGCt domain playing a critical role, as IqgD lacking RGCt fails to interact with Rac1C in a BiFC assay. In addition, we show that IqgD interacts with the actin-bundling proteins cortexillin I and II, but not simultaneously with Rac1. This mutually exclusive binding is expected because cortexillins bind to IqgD through the same GRD and RGCt domains.

Based on our results and existing knowledge about human IQGAPs and the role of Rac1 in macroendocytosis, we propose that IqgD binds active Rac1 with high affinity through its RGCt domain at macropinocytic and phagocytic cups. Similar to mammalian IQGAPs, we also show that IqgD inhibits the GTPase activity of Rac1 *in vitro*. One potential consequence of this inhibition is the protection of Rac1 from GAP-mediated inactivation, thereby prolonging spatially restricted Rac1 signaling. Such sustained Rac1 activity may be required to provide continuous input into the Scar/WAVE-Arp2/3 axis, promoting dendritic F-actin polymerization that drives persistent membrane protrusion during macroendocytic cup formation. Moreover, Rac1-bound IqgD could serve as a signaling hub that integrates Rac1 activity with F-actin binding and the recruitment of additional regulatory proteins. The function of IqgD in rear cortical structures is probably mediated by its interaction with cortexillins. We have established that the GRD is required for recruitment of IqgD to the rear cortex, where the actin-binding cortexillins are localized (38, 44), and that IqgD binds directly to cortexillins. Notably, in contrast to DGAP1 and GAPA, this interaction appears to be inhibited by Rac1 binding to IqgD. Given that the concentration of active Rac1 is lower in the posterior cortex (43), we propose that IqgD is predominantly associated with cortexillins in this region. However, the functional consequences of this interaction remain unknown and will need to be clarified in future studies. Based on the biochemical and interaction data, we propose that IqgD affects F-actin through at least two mechanisms: directly by binding filament sides via the CHD, and indirectly through its interaction with cortexillins. Consistent with this hypothesis, we find that IqgD does not alter the overall cellular F-actin content, suggesting that, unlike human IQGAP1, IqgD is unlikely to regulate actin polymerization at the filament barbed end (26, 27).

Despite its striking localization at macropinocytic and phagocytic cups, IqgD does not appear to be required for macroendocytosis. Its loss does not affect macropinosome size or the efficiency of macropinocytosis and phagocytosis from suspension. In contrast, it is critical for efficient phagocytosis of surface-bound particles, as *iqgD*^*L1*-^ cells grow almost twice as slowly as wild-type cells on bacterial lawns and display significantly reduced cell size. Notably, recent work revealed the mechanism by which macrophages internalize surface-bound particles (48). When a macrophage encounters a particle beneath its cell body, it forms an F-actin-rich circular structure around the particle at the cell-substrate interface. This structure is both adhesive and phagocytic, as it contains integrin β_2_, talin, vinculin, F-actin, Arp3, pFAK (phosphorylated focal adhesion kinase), and Rab5a (59, 60), and has therefore been termed the phagocytic adhesion ring (PAR) (48). The authors proposed a model for PAR-mediated phagocytosis in which macrophages sense a surface-attached particle through disruption of continuous waves of actin polymerization at the basal cell surface. This interference triggers PAR formation, and integrins within the PAR support inward Arp2/3-mediated actin polymerization toward the particle-substrate interface, generating the force required for particle detachment from the substrate and subsequent phagocytosis (48). Because phagocytosis is an evolutionarily conserved process, with *D. discoideum* serving as a well-established macrophage-like model (61), we hypothesize that *D. discoideum* forms a structure similar to the PAR in macrophages. Using silicone grease droplets to mimic surface-bound particles, we show that *D. discoideum* cells likewise assemble a circular F-actin-containing structure around particles at the basal cell surface. We further demonstrate that IqgD and Rac1A are components of this structure. Finally, we show that *iqgD*^*L1*-^ cells phagocytose significantly fewer surface-bound beads compared to wild-type cells. Therefore, these findings indicate that IqgD is required for efficient phagocytosis of surface-attached bacteria on bacterial lawns.

It will be interesting to investigate the composition of this ring structure in more detail in future work. Although *D. discoideum* does not express integrins (62), it is likely that other adhesion-related molecules are involved in the process because integrin-dependent tension is required for PAR formation in macrophages (48). In line with this, *iqgD*^*L1*-^ cells also have reduced cell-substrate adhesion, which probably contributes to reduced phagocytosis of surface-bound beads.

## Materials and Methods

### Plasmid vectors

In this study, vectors were used for the expression of FL and truncated variants of IqgD and other fluorescent probes in *D. discoideum* cells, for yeast two-hybrid and BiFC assays, for the expression of GST-tagged IqgD and Rac1A proteins in *E. coli*, for the expression of TwinStrep-tagged IqgD proteins in insect cells, and for the genetic inactivation of *iqgD* and *rac1A*. Detailed information is provided in the SI Appendix, SI Materials and Methods.

### Cell culture and generation of *iqgD*^*L1-*^, *iqgD*^*L1-/L2-*^, and *rac1A*^-^ cells

Cell cultivation, transfection, and gene disruption by homologous recombination were performed as previously described (63). Detailed information is provided in the SI Appendix, SI Materials and Methods.

### Quantitative real-time PCR

To compare the relative expression of the *iqgD* gene between wild-type and *iqgD*^*L1*-^ cells, qPCR was performed. Quantification was carried out using the Pfaffl method. Detailed information is provided in the SI Appendix, SI Materials and Methods.

### Yeast two-hybrid assay

The yeast two-hybrid assay was performed using the Matchmaker GAL4 Two-Hybrid System 3 (Clontech Laboratories) as previously described (39). Detailed information is provided in the SI Appendix, SI Materials and Methods.

### Protein purification and anti-IqgD antibody production

GST-tagged proteins were expressed in *E. coli* and affinity purified using glutathione-agarose 4B (Macherey-Nagel) following standard procedures. For the actin co-sedimentation assay and production of the anti-IqgD antibody, the GST tag was cleaved off from GST-IqgD_1-575. Polyclonal antibodies against IqgD were raised by immunizing female New Zealand white rabbits (Charles River) with tag-free IqgD_1-575 following standard procedures. Recombinant IqgD_1-592 and IqgD_311-1385 proteins were purified from insect cells at the EMBL Protein Expression and Purification Core Facility in Heidelberg, Germany. Detailed information is provided in the SI Appendix, SI Materials and Methods.

### Co-immunoprecipitation and GST pull-down assay

These methods were performed following standard procedures. Detailed information is provided in the SI Appendix, SI Materials and Methods.

### GAP assay

The GAP assay was performed using the GTPase-Glo Assay kit (Promega) according to the manufacturer’s instructions. Detailed information is provided in the SI Appendix, SI Materials and Methods.

### Actin co-sedimentation assay

The assay was performed as previously described (64). Detailed information is provided in the SI Appendix, SI Materials and Methods.

### F-/G-actin ratio

To determine the F-/G-actin ratio, a previously described protocol was adapted (65). Detailed information is provided in the SI Appendix, SI Materials and Methods.

### Phagocytosis

A clearance assay was used to measure phagocytosis of bacteria from suspension as previously described (66). Detailed information is provided in the SI Appendix, SI Materials and Methods.

### Cell-substrate adhesion

To assess cell-substrate adhesion, cell detachment experiments were performed as previously described (67). Detailed information is provided in the SI Appendix, SI Materials and Methods.

### Fluorescence microscopy

Confocal microscopy was performed using a Leica TCS SP8 X laser scanning confocal microscope equipped with an HC PL APO CS2 63×/1.4 oil objective, a 405 nm diode laser, and a supercontinuum excitation laser (Leica Microsystems). To analyze macropinosome size, live cells were imaged using a Dragonfly 505 spinning disc confocal microscope (Andor, Oxford Instruments) equipped with a 100×/1.45 oil objective (Nikon) and a Sona 4.2B-6 (sCMOS) camera (Andor, Oxford Instruments). Time-lapse TIRF imaging was performed using an Eclipse TI-E inverted microscope (Nikon) equipped with a TIRF Apo 100×/1.45 NA objective and an iXon3 897 EMCCD camera (Andor). Image analyses were performed using *ImageJ* (68). Cell motility was analyzed using the *ImageJ* plug-in *TrackMate* (69, 70) and Microsoft Excel with integrated DiPer macros (71). Detailed information is provided in the SI Appendix, SI Materials and Methods.

## Statistical analysis

The statistical test used for each experiment is indicated in the corresponding figure legend. All statistical analyses were performed using GraphPad Prism 5.

## Acknowledgments

This work has been supported by the Croatian Science Foundation under the project IP-2020-02-1572 to V.F.

## Author Contributions

V.F. designed research; all authors performed research; A.Č., D.P., M.Š., J.S., L.H., M.M.C, and V.F. analyzed data; and V.F. wrote the paper with contributions from other authors.

## Competing Interest Statement

The authors declare no competing interest.

## Notes

### Competing Interest Statement

The authors have declared no competing interest.

