## Supplementary Data for "IqgD is a Rac1-interacting IQGAP required for efficient growth of *Dictyostelium discoideum* on bacterial lawns"

**This PDF file includes:**

Supporting Text

Figures S1 to S7

Table S1

Legends for Movies S1-S17

SI References

**Other supporting materials for this manuscript include the following:**

Movies S1 to S17

### Supporting Text

### SI Materials and Methods

#### Plasmid vectors

The coding DNA sequence (CDS) of IqgD was amplified by PCR from cDNA prepared from vegetative cells of *Dictyostelium discoideum* laboratory strain AX2 and cloned into the pUC18 cloning vector. For expression of IqgD in *D. discoideum* cells, either unlabeled or N- and C-terminally labeled with YFP, the CDS of IqgD was amplified by PCR from pUC18_IqgD and inserted into the pDM304 (1), pDM304_N-YFP (2) or pDM304_C-YFP (this work) vectors by ligation-independent cloning (In-Fusion HD Cloning Kit, Takara Bio). For expression of YFP-tagged truncated variants of IqgD, the CDSs of IqgD_ΔCHD (Δ53-324), IqgD_ΔGRD (Δ662-1074) and IqgD_ΔRGCt (Δ1292-1385) were inserted into the pDM304_N-YFP vector by ligation-independent cloning. For co-expression of YFP-IqgD and IqgD-YFP with probes for F-actin and active Rac1 in *D. discoideum* cells, the expression cassettes containing Lifeact-mRFP and DPAKa_GBD-mRFP were excised with NgoMIV from the shuttle vectors pDM330_Lifeact-mRFP (2) and pDM330_DPAKa_GBD-mRFP (this work) and ligated into the NgoMIV site of pDM304_YFP-IqgD or pDM304_IqgD-YFP. For expression of YFP-labeled probes for active Rac1 and F-actin in *D. discoideum* cells, the previously constructed pDM304_DPAKa_GBD-YFP was used (3), and oligonucleotides encoding Lifeact were hybridized and ligated into the pDM304_N-YFP vector.

The pDM1207_GFP-mRaf1_RBD (4) was used to determine macropinosome size in wild-type and *iqgD*$^{L1-}$ cells. To determine macropinosome size in IqgD-overexpressing cells, the GFP-mRaf1_RBD-containing expression cassette was cut with NgoMIV from pDM1019_mRaf1_RBD (4) and ligated into the NgoMIV site of pDM358 (1). This vector was co-transfected with the pDM1208_IqgD vector in wild-type cells. For the pDM1208_IqgD vector, the CDS of IqgD was amplified by PCR and inserted into the pDM1208 vector (5) by ligation-independent cloning.

For the yeast two-hybrid assay, pGBKT7- and pGADT7-derived vectors were used (Matchmaker™ GAL4 Two-Hybrid System 3, Clontech). The CDS of IqgD was cloned into pGADT7 in two steps. The N-terminal half was amplified by PCR and cloned into the BglII and BamHI sites of pGADT7. Then the C-terminal half was cut from the cloning vector pUC18_IqgD and ligated into the BamHI and SacI sites of pGADT7_N-IqgD. The CDS of IqgD_GRD (662-1074) was amplified by PCR and cloned into the BamHI and SacI sites of the pGADT7 vector. The CDS of IqgD was cloned into pGBKT7 in two steps. The N-terminal half was amplified by PCR and cloned into BglII and BamHI of pGBKT7. Then the C-terminal part was amplified by PCR and cloned into the BamHI and SalI sites of pGBKT7_N-IqgD. Vector pGADT7-IqgC was previously constructed (6).

The tested GTPases of the Ras and Rho family were cloned into the pGBKT7 vector as ΔCAAX truncations. The pGBKT7 vectors for RasB(G15V/Q64L/S20N), RasG(G12V/Q61L/S17N), RasS(G12V/S17N), RapA(G14V/Q65E), Rac1A(G12V/Q61L/T17N), Rac1C(T17N), RacB(G12V), RacC(G15V/T20N) and RacG(G12V) were constructed previously (2, 6–8). Rac1B(wt)_ΔCAAX was amplified by PCR from pDGFP-Rac1B(wt) (9), cloned into the BamHI and EcoRI sites of pGBKT7, and then mutated to pGBKT7_Rac1B(G12V/T17N) by site-directed mutagenesis (Q5 site-directed mutagenesis kit, NEB). To generate pGBKT7_Rac1C(G12V) and pGBKT7_RacB(Q61L/T17N), the previously constructed pGBKT7_Rac1C(T17N) and pGBKT7_RacB(G12V) were sequentially mutated by site-directed mutagenesis. To obtain pGBKT7_RacG(T17N), pGBKT7_RacG(G12V) was mutated.

For the BiFC assay in living *D. discoideum iqgD*$^{L1-}$ cells, the previously described vectors pDM304_N-VC and pDM344_N-VN were used (2). The CDSs of full-length (FL) IqgD, IqgD_GRD (662-1074), IqgD_ΔGRD, IqgD_ΔRGCt and IqgD_311-1385 were inserted into pDM304_N-VC by ligation-independent cloning. The CDSs of the potential binding partners Rac1A, Rac1B, and Rac1C, their respective mutant forms, as well as cortexillins I and II, were cloned into pDM344_N-VN. To allow simultaneous expression of VC-IqgD and VN-Rac1 from the same vector, the expression cassettes encoding VN-Rac1A(wt/G12V/Q61L/T17N), VN-Rac1B(wt), and VN-

Rac1C(wt/G12V/T17N) were excised with NgoMIV and ligated into the NgoMIV site of pDM304_VC-IqgD. The expression cassettes encoding VN-Rac1C(Q61L), VN-CI and VN-CII were ligated into the NgoMIV site of pDM358 and these vectors were co-transfected with pDM304 encoding the respective VC-IqgD variant. To analyze the intensity of Venus signals resulting from the interaction of IqgD and Rac1 on macropinocytic cups, the BiFC vectors pDM304_VC-IqgD_VN-Rac1A/Rac1B/Rac1C(wt) were co-transfected with pDM358_mRFP-mRaf1_RBD.

For the purification of the GST-tagged FL IqgD and the truncated protein IqgD_1-575 from *Escherichia coli* cells, the CDS of IqgD was inserted into the pGEX-6P-1 vector by ligation-independent cloning and the CDS of IqgD_1-575 was amplified by PCR and ligated into the BamHI and SalI sites of pGEX-6P-1. For the GST-Rac1A pull-down assays, pGEX-5X-1_Rac1A(wt) (10), pGEX-6P-1_Rac1A(Q61L) and pGEX-6P-1_Rac1A(T17N) were used. Rac1A(Q61L) and Rac1A(T17N) were amplified by PCR from pDM344_VN-Rac1A(Q61L/T17N) and cloned into the BamHI and SalI sites of pGEX-6P-1 vector. To construct vectors for the expression and purification of the truncated IqgD_1-592 and IqgD_311-1385 proteins from insect cells, their CDSs were amplified by PCR and inserted into the pCoofy51 vector (11) by ligation-independent cloning.

The pLPBLP_iqgD$^{L1}$_KO vector was constructed to generate *iqgD*$^{L1-}$ cells. The 3′ homology region of the *iqgD*$^{L1}$ gene was amplified by PCR and cloned into the PstI and BamHI sites of the pLPBLP vector (12). The 5' homology region extending upstream into the promoter was synthesized (GenScript) and ligated into the SalI and HindIII sites of the same pLPBLP vector. To obtain *iqgD*$^{L1-}$ cells sensitive to blasticidin, the blasticidin resistance cassette was excised by Cre recombinase, transiently expressed from the pTX-NLS-cre vector (13). The pLPBLP_iqgD$^{L2}$_KO vector was constructed to generate *iqgD*$^{L1-/L2-}$ cells. The 5' and 3′ homology regions of the *iqgD*$^{L2}$ gene were amplified by PCR and cloned into the SalI and HindIII, and PstI and BamHI sites of the pLPBLP vector, respectively. The pDM1490 vector containing a hygromycin resistance cassette between the 5′ and 3′ homology regions of the *rac1A* gene (5) (a gift from Professor Arjan Kortholt, University of Groningen) was used to generate *rac1A*$^{-}$ cells.

Supporting Table S1 lists oligonucleotides used in this work.

### Cell culture

*D. discoideum* axenic strain AX2 was used as the parental strain to generate *iqgD*$^{-}$ and *rac1A*$^{-}$ cells. *Dgap1*$^{-}$*/gapA*$^{-}$ cells were generated previously (14). Cells were cultivated at 22 °C, either attached to cell culture dishes or shaken in suspension at 150 rpm in HL5 medium without glucose (Formedium), supplemented with 18 g/l maltose (Sigma-Aldrich), 50 µg/ml ampicillin (Sigma-Aldrich), and 40 µg/ml streptomycin (Fluka). Transfection by electroporation and clone selection were performed as previously described (13).

To analyze the growth rate in shaken suspension, $5 \times 10^4$ mid-log phase cells were inoculated into 20 ml of medium in 100 ml-flasks. Cell concentration was determined daily for 5 days using a CellDrop counter (DeNovix). To analyze growth on bacterial lawns of *Klebsiella aerogenes* or *E. coli* B/2, cells were plated with a dense suspension of bacteria on SM agar as for clonal selection. Plaques were imaged after 4 days on lawns of *K. aerogenes* and after 5 days on lawns of *E. coli* B/2. Plaque diameters were measured using *ImageJ* (15).

### Generation of *iqgD*$^{L1-}$, *iqgD*$^{L1-/L2-}$, and *rac1A*$^{-}$ cells

To generate *iqgD*$^{L1-}$ cells, wild-type AX2 cells were transfected with 35 µg of BamHI/SalI-linearized and dephosphorylated pLPBLP_iqgD$^{L1}$_KO vector. Transfectants were selected in HL5 medium with 10 µg/ml blasticidin S (Sigma-Aldrich) for 14 days. After clonal selection, initial screening for *iqgD*$^{L1}$ disruption was performed by PCR. The gDNA of potential clones was digested with ClaI and analyzed by Southern blot using a bsr probe complementary to the CDS of blasticidin S deaminase. The membrane was then stripped and rehybridized with the R probe complementary to the *iqgD* gene. Inactivation of the *iqgD*$^{L1}$ gene (clone IW047) was also confirmed by western blot using anti-IqgD antibody. The *iqgD* gene was disrupted in the same way in AX2-214 cells newly obtained from

Professor Annette Müller-Taubenberger (Ludwig Maximilian University of Munich), and successful inactivation of the gene was confirmed by PCR and western blot.

To generate *iqgD*$^{L1-/L2-}$ cells, the bsr cassette was removed from the *iqgD*$^{L1}$ locus. *IqgD*$^{L1-}$ cells were transfected with 35 µg of the pTX-NLS-Cre vector and cultivated in HL5 medium with 10 µg/ml G418 for 8 days. After clonal selection on a bacterial lawn, cells sensitive to both blasticidin S and G418 were selected, and excision of the bsr cassette was confirmed by PCR. The *iqgD*$^{L1-}$ cells with the bsr cassette removed were transfected with 35 µg of BamHI/SalI-linearized and dephosphorylated pLPBLP_iqgD$^{L2}$_KO vector. Transfectants were selected as previously described. Disruption of *iqgD*$^{L2}$ (clone IW052) was confirmed by PCR and sequencing.

*Rac1A*$^{-}$ cells were generated by transfecting wild-type AX2 cells with 10 µg of PvuII-digested and dephosphorylated pDM1490 containing the 5′ and 3′ homology regions of the *rac1A* gene. Clones were selected in 96-well plates in HL5 medium with 50 µg/ml hygromycin B (Sigma-Aldrich). Transfectants were diluted and plated at a density of approximately 0.3 cell/well. Initial screening for *rac1A* gene disruption was performed by PCR. The gDNA of positive clones was digested with ClaI and analyzed by Southern blot using a hygr probe specific for the CDS of hygromycin phosphotransferase. The membrane was then stripped and rehybridized with the L probe complementary to the *rac1A* gene. We named this *rac1A*$^{-}$ clone IW061.

A PCR DIG Probe Synthesis Kit (Roche) was used to label DNA probes for Southern blot with digoxigenin, and the hybridized DNA bands were visualized using a DIG Luminescent Detection Kit (Roche) on the Alliance Q9 Mini Chemiluminescence Imaging System (Uvitec).

#### Quantitative real-time PCR

To compare the relative expression of the *iqgD* gene between wild-type and *iqgD*$^{L1-}$ cells, total RNA was extracted using the RNeasy Mini Kit (Qiagen) according to the manufacturer's instructions and reverse transcribed into cDNA using the High Capacity RNA-to-cDNA Kit (Applied Biosystems). The resulting cDNA was used as a template in qPCR with Power SYBR Green PCR Master Mix (Applied Biosystems) on the ABI 7300 Real-Time PCR system (Applied Biosystems). To quantify *iqgD* gene expression in *iqgD*$^{L1-}$ cells relative to wild-type cells, the Pfaffl method was used, which accounts for the amplification efficiency of each gene (16). *IqgD* gene expression was normalized to the expression level of the housekeeping gene *gpdA*.

#### Yeast two-hybrid assay

To investigate direct protein interactions using the yeast two-hybrid assay, we used the Matchmaker GAL4 Two-Hybrid System 3 (Clontech Laboratories). The *Saccharomyces cerevisiae* strain AH109 was co-transfected with the pGBKT7- and pGADT7-derived plasmid pair. Yeast growth on selection plates lacking leucine and tryptophan (-LW) confirmed efficient co-transfection. Direct protein interactions were analyzed on selection plates lacking leucine, tryptophan, and histidine, with 0.5 mM 3-amino-1,2,4-triazole (3-AT).

#### Purification of GST-IqgD_1-575 and anti-IqgD antibody production

*E. coli* Rosetta 2 cells (Novagen) were transformed with the pGEX-6P-1-IqgD_1-575 vector, and expression of recombinant GST-tagged IqgD_1-575 protein was induced overnight with 0.75 mM isopropyl-β-D-thiogalactoside (IPTG) at 21 °C and 150 rpm. The bacteria were harvested and lysed by ultrasonication in lysis buffer (30 mM Hepes pH 7.4, 150 mM NaCl, 2 mM EDTA, 1 mM DTT, 5% glycerol, 5 mM benzamidine hydrochloride, 0.5 mM 4-(2-aminoethyl)benzenesulfonyl fluoride hydrochloride, and 2 units/ml Benzonase). The fusion protein was purified from bacterial extracts by affinity chromatography using glutathione-agarose 4B (Macherey-Nagel) following standard procedures. The GST tag was cleaved off by PreScission protease (GE Healthcare) and removed by absorption on fresh glutathione-agarose. Proteins in the flow-through were then separated by size-exclusion chromatography using a HiLoad 26/600 Superdex 200 column controlled by an Äkta Purifier System. Protein concentration was determined by absorption spectroscopy using extinction coefficients derived from the amino acid sequence. Polyclonal antibodies against IqgD were raised by immunizing female New Zealand white rabbits (Charles River) with tag-free IqgD_1-575 following standard procedures.

**Statement on care and use of animals**

The immunization of rabbits for the generation of polyclonal antibodies was conducted in accordance with national guidelines for the care and maintenance of laboratory animals and approved by the Hannover Medical School Institutional Animal Care Facility and the Lower Saxony State Office for Consumer Protection and Food Safety under application number 22-00236 to J.F.

**Purification of recombinant IqgD_1-592 and IqgD_311-1385 from insect cells**

Purification of recombinant proteins was performed at EMBL Protein Expression and Purification Core Facility in Heidelberg, Germany. The pCoofy51 plasmids carrying the N-terminal TwinStrep-tagged IqgD_1-592 and IqgD_311-1385 genes were used for transposition into *E. coli* DH10EMBacY cells (Geneva Biotech). Insertion of the i*qgD* CDSs into the bacmid DNA was verified by PCR. The purified bacmid DNA was transfected into Sf9 insect cells using XtremeGENE HP (Sigma-Aldrich) DNA transfection reagent. After three days at 27 °C, YFP-positive cells were observed using fluorescence microscopy and the V0 baculoviruses were harvested. To amplify the recombinant baculoviruses, 25 ml of Sf9 cells at a density of 1 × $10^6$ cells/ml were infected with 3 ml of the V0. The Sf9 suspension cultures were incubated for 3 days at 27 °C, after which the V1 baculovirus stocks were harvested. For large scale expression, 4 × 1 l of Sf21 suspension cultures in Sf-900 III SFM medium (Thermo Fischer Scientific) at a density of 1 × $10^6$ cells/ml were infected with 3.3 ml of the PKH V1 baculoviruses (1:300 dilution). The infected Sf21 cultures were incubated at 27 °C. After 3 days, the cultures were harvested by centrifugation (20 min, 550 × g, 4 °C) and the cell pellets were flash-frozen and stored at -80 °C. For protein purification, the cell pellets were thawed and resuspended in cold lysis buffer (50 mM Tris-HCl pH 8.0, 500 mM NaCl, 1 mM DTT, 10% glycerol, 2 mM $MgCl_2$, SmNuclease and cOmplete EDTA-free Protease Inhibitors, Roche). All further purification steps were performed at 4-8 °C. Cells were lysed by 5 passages through a microfluidizer device, followed by centrifugation (30 min, 140,000 × g, 4 °C). The cleared lysate was loaded onto a 5 ml Strep-Tactin Superflow High Capacity column (IBA) preequilibrated with 50 mM Tris-HCl pH 8.0, 500 mM NaCl, 1 mM DTT and 10% glycerol. After loading, the Strep-Tactin column was washed with equilibration buffer and eluted with equilibration buffer supplemented with 5 mM Desthiobiotin. After SDS-PAGE analysis, elution fractions containing the respective IqgD proteins were pooled, concentrated to ~ 1 ml and injected onto a Superdex 200 Increase 10/300 size exclusion chromatography column (Cytiva) preequilibrated with 50 mM Tris pH 8.0, 300 mM NaCl, 1 mM DTT, 2 mM $MgCl_2$ and 10% glycerol. The IqgD-containing elution fractions were pooled and concentrated to ~ 1-2 mg/ml. The final IqgD samples were aliquoted, flash-frozen in liquid nitrogen and stored at -80 °C until usage. Identity and intactness of the purified IqgD_1-592 and IqgD_311-1385 proteins were verified by mass spectrometry (performed by the EMBL Proteomics Core Facility). The oligomerization state of both proteins was assessed by mass photometry using the Refeyn TwoMP instrument. Both IqgD_1-592 and IqgD_311-1385 were shown to be monomeric, although for IqgD_311-1385 some additional species with higher and lower molecular weight could be observed as well.

**Co-immunoprecipitation**

For co-immunoprecipitation of actin and cortexillins I and II with endogenous IqgD, a polyclonal anti-IqgD antibody was bound to Protein A–Sepharose CL-4B (GE Healthcare), followed by overnight incubation of the beads with lysates from wild-type and IqgD-overexpressing cells. For the negative control, a nonspecific antibody was used. After elution of the protein complexes captured by the antibody, the samples were analyzed by immunoblotting using anti-actin mAb 224-236-1 (17), anti-cortexillin I mAb 241-438-1 (18), and anti-cortexillin II mAb 232-238-10 (18) antibodies.

### GST pull-down assays

*E. coli* Rosetta 2 cells (Novagen) were transformed with the pGEX-5X-1_Rac1A(wt) and pGEX-6P-1_Rac1A(Q61L/T17N) vectors, and expression of GST-Rac1A(wt/Q61L/T17N) was induced overnight with 0.25 mM IPTG at 25 °C and 150 rpm. The recombinant proteins were affinity-purified from bacterial lysate using glutathione-agarose (Thermo Scientific) according to the manufacturer's instructions. Purified GST-Rac1A(wt) protein was loaded with GDP or GTPγS as previously described (10). To examine whether IqgD exists in a complex with Rac1A, lysates of wild-type and IqgD-overexpressing cells were incubated with immobilized GST-Rac1A(wt) for 2 hours at 4 °C as previously described (3). To test IqgD binding to mutant variants of Rac1A, cell lysates were incubated with immobilized GST-Rac1A(Q61L/T17N) overnight at 4 °C. To test direct binding of IqgD to Rac1A, 5 µg of truncated IqgD_311-1385 purified from insect cells was incubated with GDP- and GTPγS-loaded GST-Rac1A for 2 hours at 4 °C. IqgD binding was analyzed by immunoblotting with polyclonal anti-IqgD antibody. To investigate if cortexilins bind to IqgD bound to Rac1A, lysate of *dgap1⁻/gapA⁻* cells was incubated with IqgD_311-1385 bound to GST-Rac1A overnight at 4 °C. Cortexilin binding was analyzed by immunoblotting with anti-cortexillin I mAb 241-438-1 (18) and anti-cortexillin II mAb 232-238-10 (18) antibodies. The lysis buffer used in pull-down assays contained: 30 mM Tris-HCl pH 7.5, 40 mM NaCl, 1 mM EGTA pH 7.3, 2 mM DTT, 5 mM $MgCl_2$, 0.5% n-octylpolyoxyethylene, 1 mM PMSF, 5 mM benzamidine, and cOmplete EDTA-free Protease Inhibitors Cocktail (Roche). In all pull-down assays, GST pull-down was used as a negative control.

### GAP assay

The GAP assay was performed using the GTPase-Glo Assay kit (Promega) according to the manufacturer's protocol for 384-well plates. The GTPase activity of 2 µM human Rac1(wt) (ab89246, Abcam) was measured in the absence and presence of purified IqgD_311-1385 (2 µM and 4 µM) in GAP buffer containing 5 µM GTP. Luminescence was measured with a Spark multimode plate reader (Tecan).

### Actin co-sedimentation assay

$Ca^{2+}$-ATP G-actin was purified from rabbit skeletal muscle as previously described (19) and stored on ice in G-buffer (5 mM Tris-HCl pH 8.0, 0.2 mM $CaCl_2$, 0.5 mM DTT, 0.2 mM ATP). IqgD_1-575 and IqgD_1-592 were purified from bacterial and insect cells, respectively, as described above. For co-sedimentation assays, IqgD_1-575 and IqgD_1-592 were clarified by centrifugation at 150,000 × g for 45 min at 4 °C using an Optima MAX-XP ultracentrifuge (Beckman Coulter). Next, 30 µM or 4 µM G-actin was polymerized for 2 h at room temperature in the absence or presence of 4 µM IqgD_1-592 or 4 µM IqgD_1-575 in actin polymerization buffer (10 mM imidazole pH 7.0, 2 mM $MgCl_2$, 50 mM KCl, 1 mM ATP). For high-speed co-sedimentation assays, samples were centrifuged under the same conditions as above. For low-speed co-sedimentation assays, samples were centrifuged at 17,000 × g for 30 min at 4 °C in a PICO17 centrifuge (Heraeus). The supernatants were then carefully separated from the pellets and adjusted to the same volume with SDS sample buffer. Proteins were resolved by SDS-PAGE and visualized by Coomassie Blue staining. Protein content in pellet and supernatant fractions was determined by densitometry.

### F-/G-actin ratio

To determine the F-/G-actin ratio, a previously described protocol was adapted (20). Cells grown overnight in shaken suspension to mid-log phase were collected at $2 \times 10^7$ cells, washed in cold wash buffer (10 mM Tris-HCl pH 7.5, 50 mM KCl, 1 mM EGTA, 5 mM $MgCl_2$, 0.5 mM DTT, 1 mM PMSF, 1 mM ATP), and lysed with 600 µl cold lysis buffer (wash buffer containing 5% glycerol, 1% Nonidet P-40, and cOmplete EDTA-free Protease Inhibitors, Roche). A 200 µl aliquot of lysate was collected to measure protein concentration. The remaining 400 µl of lysate was used to separate F- and G-actin by centrifugation in low-protein binding tubes (LoBind Tubes, Eppendorf). Lysate was centrifuged at 30,000 × g for 2 hours at 4 °C. The supernatant was collected, and the pellet was resuspended in 400 µl lysis buffer. The F-actin fraction was sonicated three times for 30 s at an amplitude of 60 (Ultrasonic Processor, Cole-Parmer). One microgram of protein from each fraction was separated by SDS-PAGE and immunoblotted with anti-actin mAb 224-236-1 (17). The detected bands were analyzed by densitometry using *ImageJ* (15).

#### Phagocytosis

A clearance assay was used to measure phagocytosis of bacteria from suspension (2). Mid-log phase cells grown overnight in shaken suspension were washed with ice-cold phosphate buffer, adjusted to a concentration of $2 \times 10^6$ cells/ml in 10 ml of phosphate buffer, and shaken for 30 min at 150 rpm. An equal volume of *E. coli* DH5α or B/2 suspension with an optical density at 600 nm ($OD_{600}$) of approximately 1.4 in phosphate buffer was added to the cells. Immediately and every hour for 6 hours, 1.2 ml samples were withdrawn, supplemented with Na-azide to a final concentration of 15 mM, and vortexed for 1 min to remove surface-bound bacteria. After pelleting the cells by centrifugation for 1 min at 700 × g, the $OD_{600}$ of the supernatant containing bacteria was measured. All experiments were corrected for protein content and normalized to the initial $OD_{600}$ of wild-type cells.

#### Cell-substrate adhesion

To assess cell-substrate adhesion, cell detachment experiments were performed as previously described (8). Subconfluent cells were plated on a 6 cm Petri dish in HL5 medium without selection antibiotics at a concentration of $1 \times 10^5$ cells/ml one day before the experiment. The next day, plates were shaken on an orbital shaker in fresh HL5 medium. Detached and attached cells were collected, and cell concentration was measured with a CellDrop counter (DeNovix) for both attached and detached cells before shaking (0 min), and at 30 min and 60 min after shaking at 100 rpm at 22 °C. Results are presented as the percentage of detached cells out of the total number of cells.

#### Fluorescence microscopy

Confocal microscopy was performed using a Leica TCS SP8 X laser scanning confocal microscope equipped with an HC PL APO CS2 63×/1.4 oil objective, a 405 nm diode laser, and a supercontinuum excitation laser (Leica Microsystems). To examine the (co)localization of YFP- and mRFP-tagged proteins, phagocytosis of TRITC-labeled yeast particles, and BiFC signal in living *D. discoideum*, the excitation wavelengths and detection ranges used were as follows: 511 nm and 520-565 nm for YFP, 515 nm and 525-600 nm for Venus, 565 nm and 575-650 nm for TRITC, and 585 nm and 592-650 nm for mRFP. To assess cytokinesis efficiency of cells from the vegetative zone of a spreading plaque, cells were harvested with an inoculation loop, washed free of bacteria in phosphate buffer, fixed with 2% paraformaldehyde, and stained with 1 µg/ml DAPI for 30 min. The excitation wavelength and detection range used for DAPI imaging were 405 nm and 430–520 nm. For immunostaining F-actin, cells were fixed with cold methanol for 5 min at -20 °C and then stained with anti-actin mAb 224-236-1 (17) and Alexa 633-conjugated goat anti-mouse IgG secondary antibody (Invitrogen). To estimate fluid uptake, cells were incubated in the dark in HL5 medium containing 2 mg/ml TRITC-dextran. After 10 min, cells were washed twice with phosphate buffer and immediately imaged. Each cell was outlined, and TRITC fluorescence within the outlined area was measured in *ImageJ* (15). Macropinocytosis was expressed as the ratio of TRITC intensity to cell area (21). To determine the efficiency of phagocytosis of surface-bound microbeads, a previously described protocol was adapted (22). Glass coverslips were incubated for 30 min in poly-L-lysine (0.01%, Sigma-Aldrich). After washing the coverslips three times with water, a suspension of $3.6 \times 10^8$ 1 µm microbeads (Fluoresbrite® YG Microspheres, Polysciences) in 350 µl phosphate buffer was added to the coverslips. After 20 min of incubation, an equal volume of 9% glutaraldehyde (Sigma-Aldrich) in phosphate buffer was added to the microbead solution. Following 20 min of incubation and washing the coverslips five times with water, 5 mg/ml sodium borohydride (Sigma-Aldrich) was added and incubated for 5 min to quench the glutaraldehyde. After washing the coverslips five times with phosphate buffer, a suspension of cells was added to the coverslips and incubated for 30 min at 22 °C. Cells were then fixed with cold methanol for 5 min at -20 °C. The next day, cells were imaged in *xyz* mode and analyzed in 3D to determine how many microbeads they had ingested. The excitation wavelength and detection range used for microbeads were 458 nm and 469–495 nm, respectively. For the latrunculin assay, Latrunculin A (Sigma Aldrich) at a final concentration of 5 µM was added to YFP-IqgD-expressing cells, and cells were imaged at the indicated times. After 20 min, cells were washed with phosphate buffer and further imaged. To analyze macropinosome size, live cells expressing the GFP-mRaf1_RBD probe were imaged in *xyzt* mode using a Dragonfly spinning disc confocal microscope (Andor, Oxford Instruments) with a 100×/1.45 oil objective (Nikon) and a Sona 4.2B-6 (sCMOS) camera (Andor, Oxford Instruments). The excitation wavelength was 488 nm, and the detection bandpass filter 521/38 was used. Macropinosome size was determined by measuring the length of the major axis upon macropinosome closure using *ImageJ* (15). To analyze random migration, cells were plated at low density in HL5 medium on a 35 mm glass-bottomed dish (MatTek) 1 hour before the experiment. Cells were imaged every 10 s for 1 hour using a dark-field microscope (Axiovert 135, Zeiss) with a 5× objective (23). Acquired images were processed in *ImageJ* (15), and single-cell trajectories were detected using *ImageJ* plug-in *TrackMate* (24, 25). Trajectory data were imported into Microsoft Excel with integrated DiPer macros, which were used to calculate cell speed and directional persistence (26). To assess the localization of IqgD at ventral actin dots on the substrate, *iqgD*$^{L1-}$ cells co-expressing YFP-IqgD and Lifeact-mRFP were seeded in HL5 medium onto 35 mm glass-bottomed dishes (Ibidi) at least 1 hour before imaging. Time-lapse TIRF imaging was performed using an Eclipse TI-E inverted microscope (Nikon) equipped with a TIRF Apo 100×/1.45 NA objective and an iXon3 897 EMCCD camera (Andor). Images were recorded at 5 s intervals for at least 5 min using the 488-nm and 561-nm laser lines.

**Statistical analysis**

The statistical test used for each experiment is indicated in the corresponding figure legend. All statistical analyses were performed using GraphPad Prism 5.

### Supporting Figures

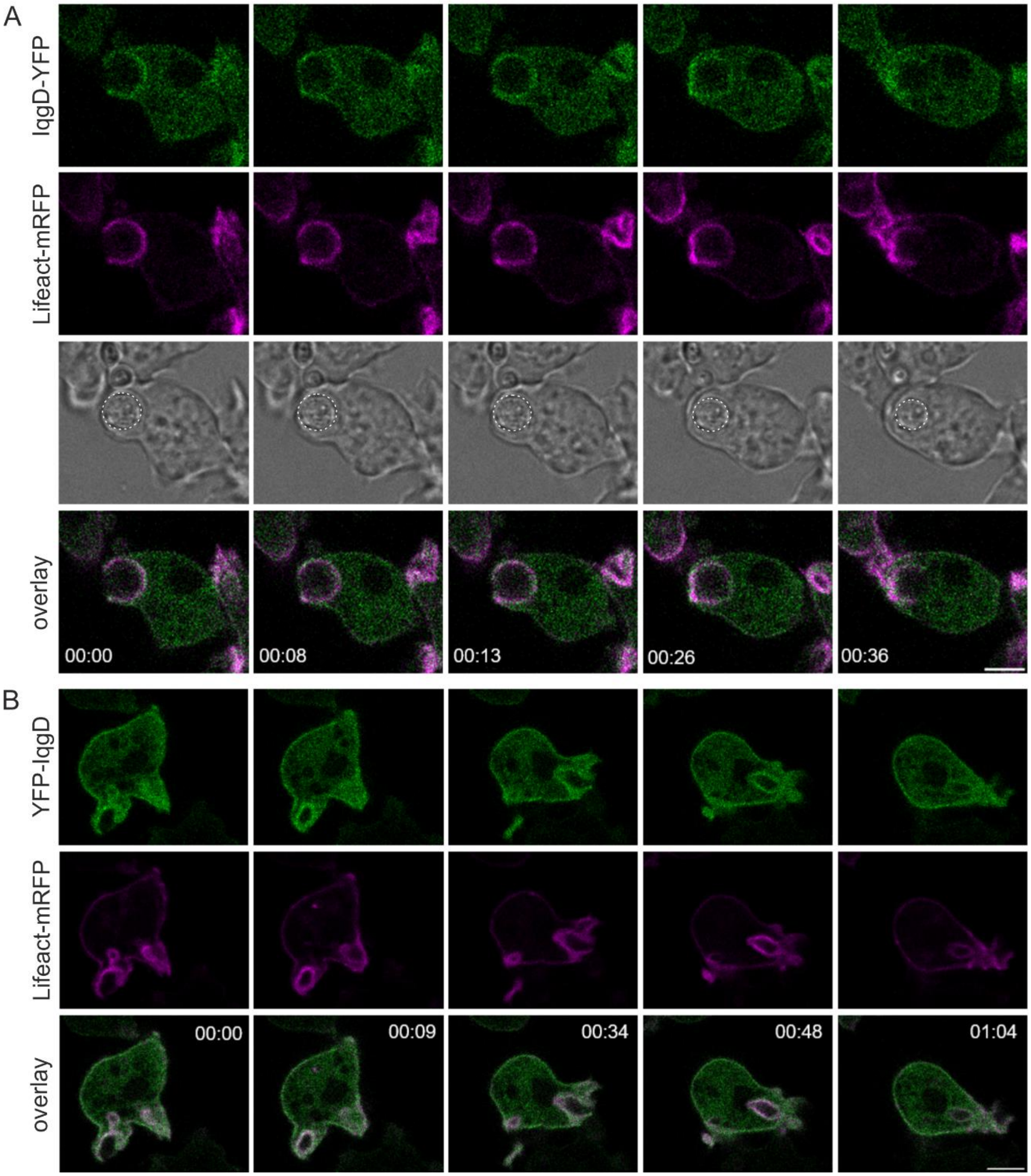


**Figure S1**. **IqgD is colocalized with the F-actin marker Lifeact-mRFP during random movement, macropinocytosis, and phagocytosis.** (A) Image sequence of a growth-phase AX2 cell ectopically expressing IqgD-YFP and Lifeact-mRFP during phagocytosis of a yeast particle. Dashes in the transmission channel encircle the yeast particle. (B) Image sequence of a growth-phase AX2 cell ectopically expressing YFP-IqgD and Lifeact-mRFP during random movement and macropinocytosis. Time is given in min:sec format. Scale bars: 5 µm. A and B correspond to Movies S5 and S6.

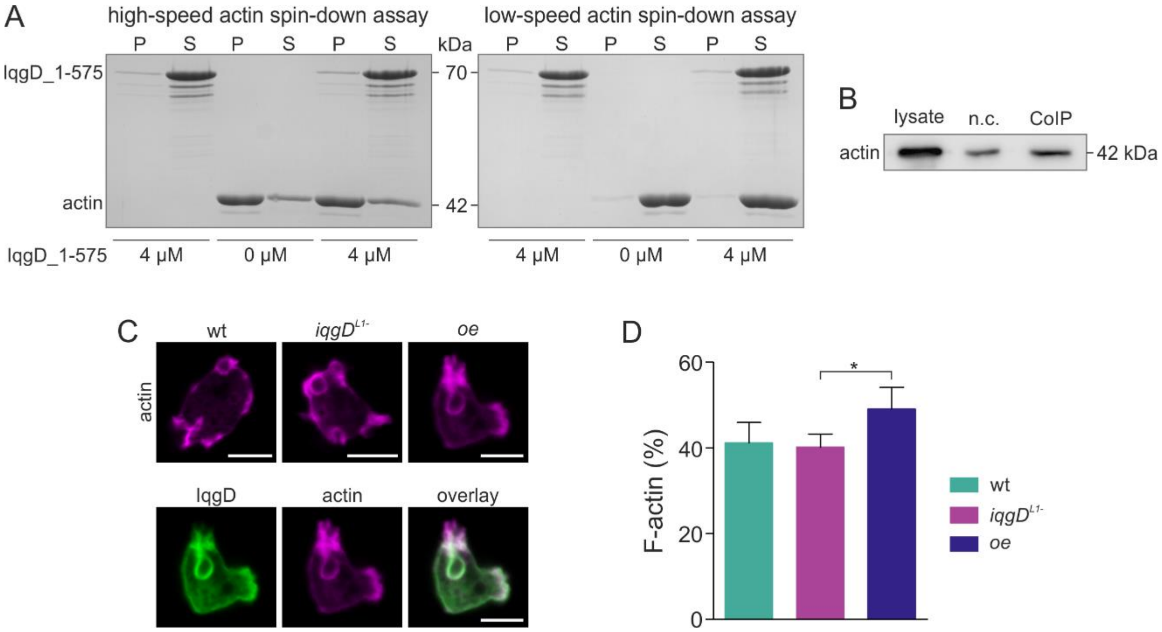


**Figure S2. (A) IqgD_1-575 produced in *E. coli* does not bind F-actin.** The CBB-stained PAGs from the high-speed (*left*) and low-speed (*right*) actin spin-down assays show that IqgD_1-575 produced in bacterial cells does not co-sediment with F-actin. *P* and *S* indicate pellet and supernatant, respectively. **(B) Endogenous actin co-immunoprecipitates with endogenous IqgD.** Anti-actin blot of co-immunoprecipitation of actin with endogenous IqgD (*CoIP*). Co-immunoprecipitation with non-specific IgG was performed as a negative control (*n.c.*). **(C) Wild-type (wt), IqgD-deficient (*iqgD$^{L1-}$*), and YFP-IqgD-overexpressing (*oe*) cells show similar actin distribution.** Representative images of wt, *iqgD$^{L1-}$*, and *oe* cells immunostained with anti-actin antibody (*upper panel*), and separate channels showing IqgD and actin in the *oe* cell from the upper panel (*lower panel*). **(D) Depletion of IqgD does not affect the amount of F-actin in the cell.** Statistical analysis was performed using the Kruskal-Wallis test followed by Dunn's multiple comparison test for pairwise comparisons. The significance level was set at 5%. * $P < 0.05$.

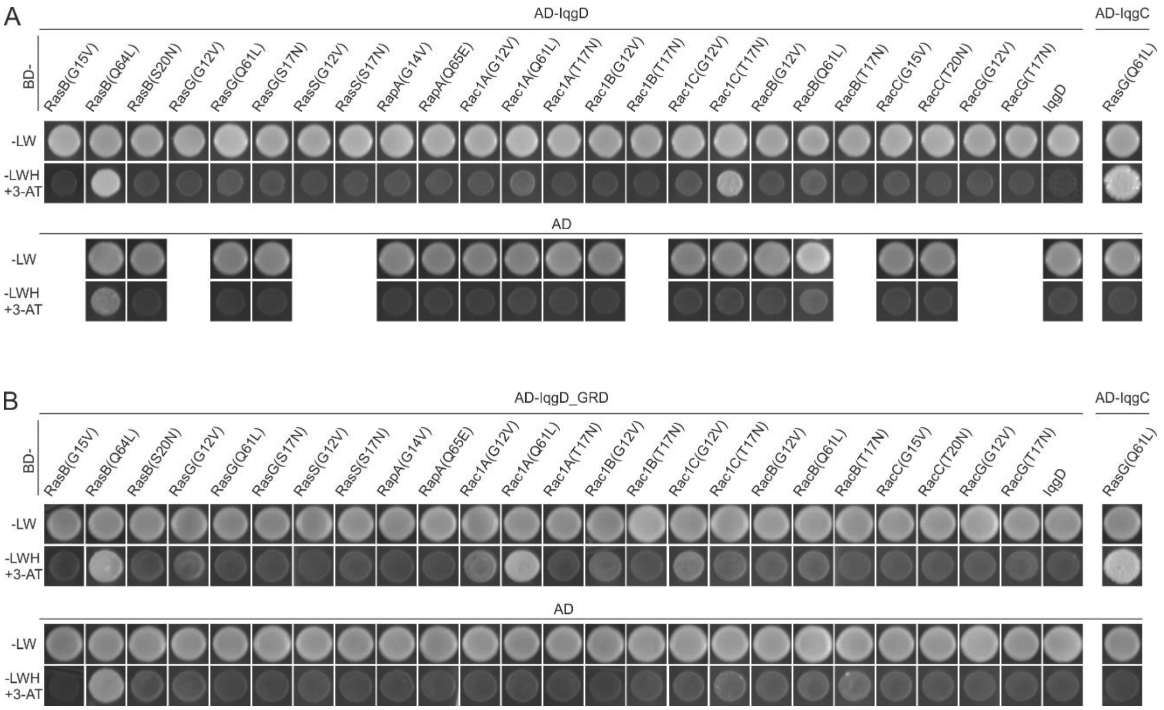


**Figure S3. IqgD interacts with Rac1 GTPases in the yeast two-hybrid assay.** (A) Full-length IqgD and (B) IqgD_GRD were fused to the Gal4 activation domain (AD) and tested GTPases to the Gal4 DNA-binding domain (BD). Successful transfection with both expression vectors was confirmed by growth of yeast transformants on selective plates lacking leucine and tryptophan (-LW). Interactions between the tested proteins were examined by growth on plates lacking leucine, tryptophan and histidine (-LWH) in the presence of 0.5 mM 3-amino-1,2,4-triazole (3-AT). The interaction between AD-IqgC and BD-RasG(Q61L) was used as a positive control. The interaction was declared positive after comparison with the growth of yeast transfected with the vector for the corresponding GTPase and the empty pGADT7 vector, i.e. with the corresponding negative control.

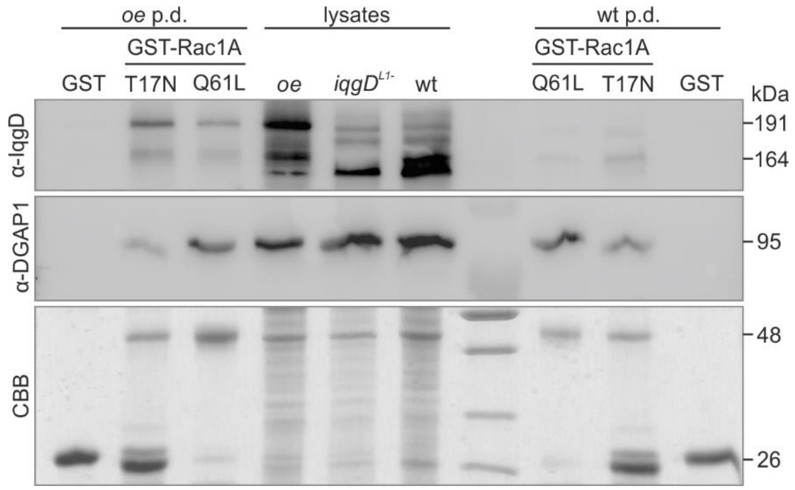


**Figure S4. Exogenous YFP-IqgD and endogenous IqgD from cell lysates bind with higher affinity to inactive Rac1A.** The anti-IqgD blot (*upper panel*) of GST-Rac1A pull-downs with lysates of wild-type cells expressing YFP-IqgD (*oe p.d., left*) and wild-type cells (*wt p.d., right*) shows that both exogenous YFP-IqgD (191 kDa) and endogenous IqgD (164 kDa) bind more strongly to constitutively inactive (T17N) than to constitutively active (Q61L) Rac1A. The anti-DGAP1 blot (*middle panel*) was used as a control, showing that, in contrast to IqgD, endogenous DGAP1 binds more strongly to constitutively active Rac1A (10). GST pull-downs were performed as a negative control. The CBB-stained PAG of GST proteins used in the pull-down assays is shown (*lower panel*).

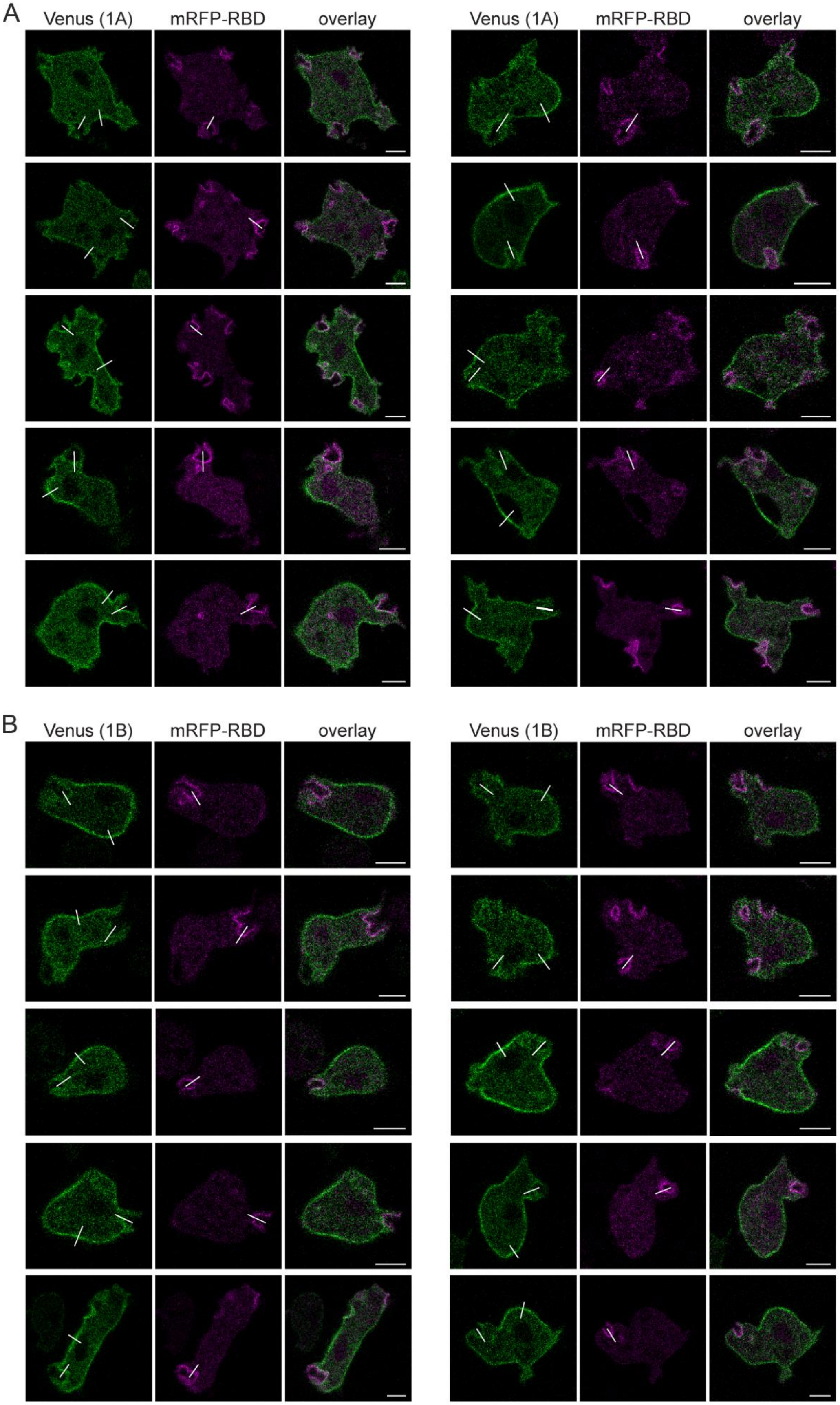
A
Venus (1A)
mRFP-RBD
overlay
Venus (1A)
mRFP-RBD
overlay
B
Venus (1B)
mRFP-RBD
overlay
Venus (1B)
mRFP-RBD
overlay

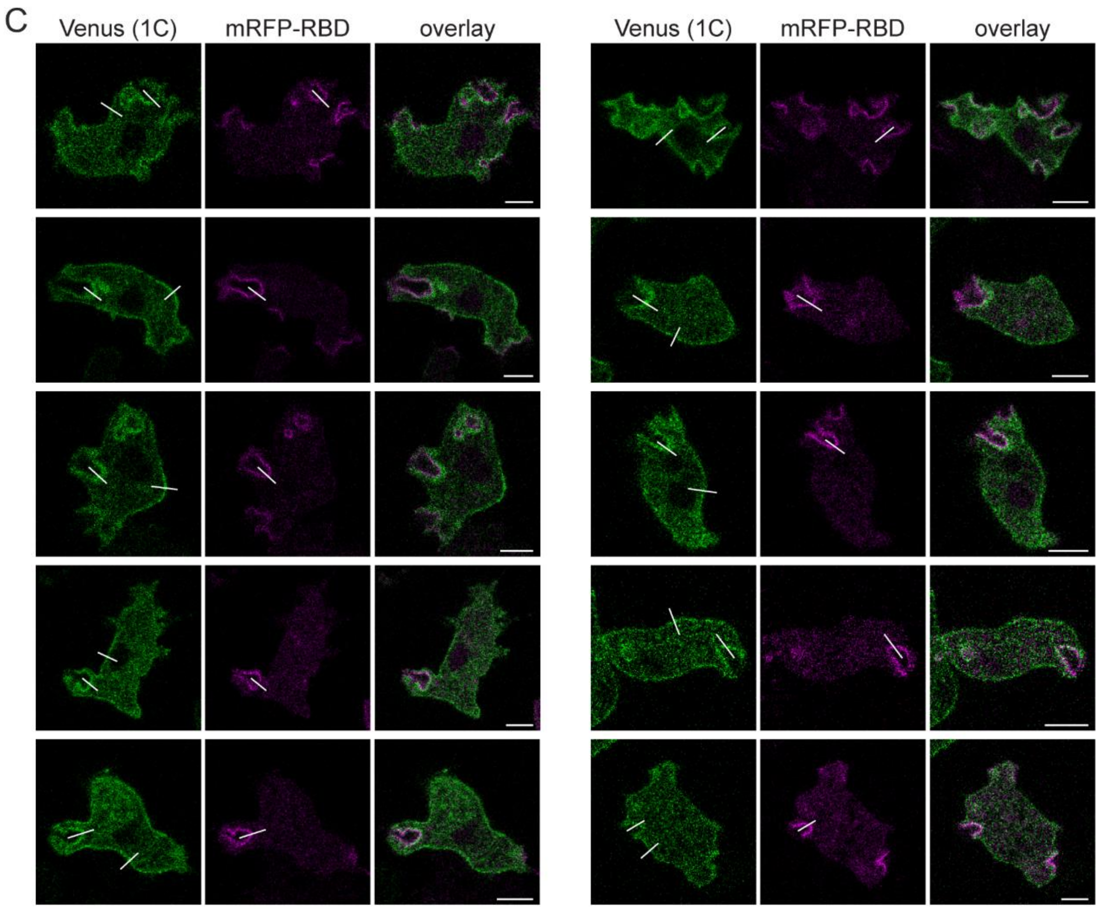


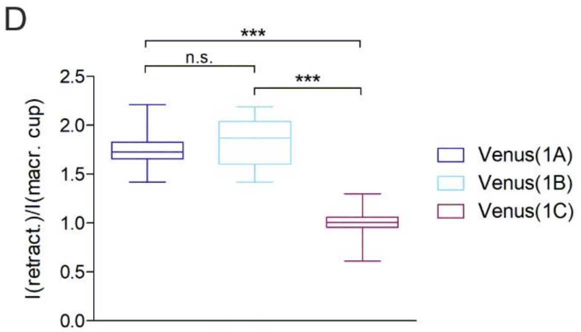


**Figure S5. Rac1 GTPases show differential binding to the IqgD population in the posterior retractive parts of the cell cortex and to the IqgD population at the macropinocytic cups.** Images of 10 *iqgD*$^{L1-}$ cells expressing VC-IqgD and VN-Rac1A(wt) (*Venus (1A)*) (A), VN-Rac1B(wt) (*Venus (1B)*) (B), or VN-Rac1C(wt) (*Venus (1C)*) (C), with mRFP-RBD. The lines represent the approximate positions where the intensities of the Venus signal were measured. Scale bars: 5 µm. D) The ratios of Venus intensity in the retracting region, *I(retract.)*, and at the base of the macropinocytic cup, *I(macr. cup)*, were determined for all cells in (A) – (C) and were as follows: 1.754 ± 0.210 for VC-IqgD and VN-Rac1A; 1.835 ± 0.264 for VC-IqgD and VN-Rac1B; and 0.994 ± 0.175 for VC-IqgD and VN-Rac1C (mean ± SD). Statistical analysis was performed using One-way ANOVA followed by Tukey's Multiple Comparison Test. The significance level was set at 5%. n.s. – not significant, *** $P < 0.001$.

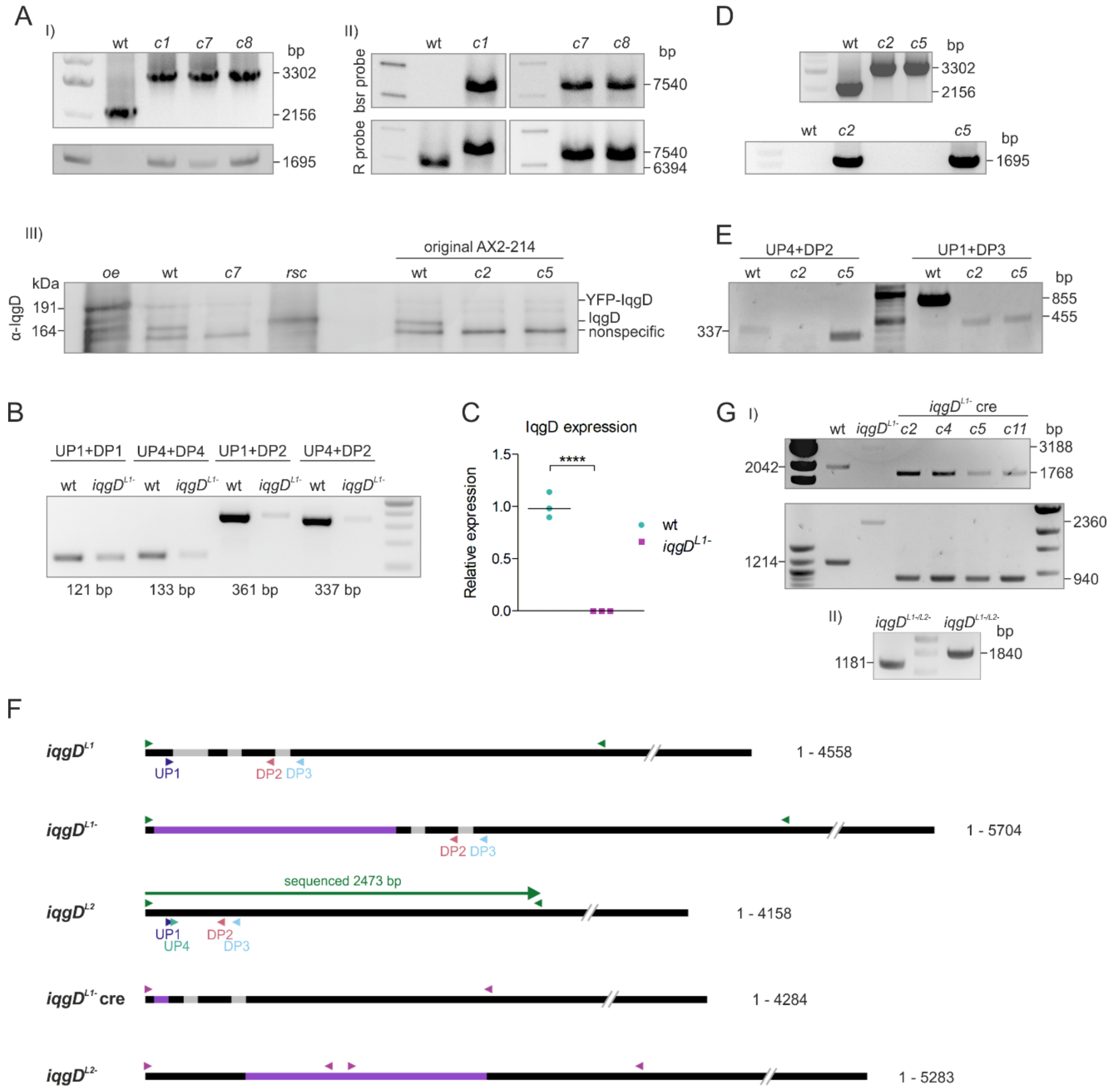


**Figure S6. The genome of AX2-214 cells contains two loci encoding IqgD.** (A) Inactivation of the *iqgD* gene on chromosome 2 (DDB_G0275731) by homologous recombination was confirmed by (I) PCR, (II) Southern and (III) western blot. (I) Two PCRs were performed to detect the integration of a blasticidin resistance (bsr) cassette at the target locus. One PCR was performed with a primer pair that yielded PCR products of different sizes for wild-type (wt, 2156 bp) and knockout (clones *c1*, *c7* and *c8*, 3302 bp) cells (*upper gel*). The other PCR was performed with a primer complementary to a bsr cassette, while the other primer is complementary to the gene sequence downstream of the 3' homology region. Therefore, the PCR product (1695 bp) can only be obtained from knockout clones (*lower gel*). (II) The Southern blot was performed with two probes, one complementary to the bsr cassette (bsr probe, *upper membrane*) and the other complementary to the gene sequence downstream of the 3' homology region (R probe, *lower membrane*). The bsr probe detects a band only in the knockout clones (*c1*, *c7* and *c8*, 7540 bp), while the R probe hybridizes with bands of different lengths in wild-type (wt, 6394 bp) and knockout (*c1*, *c7* and *c8*, 7540 bp) cells. (III) Anti-IqgD blot of lysates from wild-type cells expressing exogenous YFP-IqgD (191 kDa, *oe*), wild-type cells (wt), knockout clone 7 (*c7*) and knockout clone 7 expressing exogenous IqgD without tag (164 kDa, *rsc*) (*4 lanes on the left side of the membrane*). Three lanes on the right side of the membrane are lysates of newly obtained wild-type AX2-214 cells (*original AX2-214*) and corresponding *iqgD* knockout clones (*c2* and *c5*) obtained by homologous recombination. The band labeled „nonspecific" was initially mistaken for IqgD. To clarify the confusion, we expressed IqgD without tag in knockout cells (*see rsc*). (B) End-point PCR with different primer pairs designed for qPCR yielded extremely less PCR products from knockout (*iqgD*$^{L1-}$) compared to wild-type (wt) cells. (C) qPCR with UP1 and DP1 primers (see (B)) confirmed several hundred-fold lower expression of the IqgD transcript in *iqgD*$^{L1-}$ cells compared to wt (n = 3, median). (D) Inactivation of *iqgD*$^{L1}$ by homologous recombination in freshly obtained AX2-214 cells was confirmed by PCR and western blot (see (A), *3 lanes on the right side of the membrane*). Two PCRs were performed as described in (A), and showed successful inactivation of *iqgD*$^{L1}$ in clones *c2* and *c5*. (E) The presence of *iqgD*$^{L2}$ in newly obtained AX2-214 cells was confirmed by two PCRs. One PCR was performed with a primer pair specific for the *iqgD*$^{L2}$ locus and yielded PCR product (337 bp) for both wild-type and *iqgD*$^{L1-}$ clones *c2* and *c5* (*left of the DNA standard on the gel*). The other PCR was performed with primers that yielded products of different lengths from both loci (855 bp from *iqgD*$^{L1}$ and 455 bp from *iqgD*$^{L2}$). Therefore, two PCR products were obtained in the wt sample, with the shorter product only faintly visible, while only the shorter PCR product was obtained in *iqgD*$^{L1-}$ clones *c2* and *c5* (*to the right of the DNA standard on the gel*). (F) Schematic representation of the *iqgD* gene at the known locus on chromosome 2 before (***iqgD***$^{L1}$) and after (***iqgD***$^{L1-}$) homologous recombination; the *iqgD* gene at an unknown genomic position before (***iqgD***$^{L2}$) and after (***iqgD***$^{L2-}$) homologous recombination; and *iqgD*$^{L1-}$ after excision of the bsr cassette with Cre recombinase (***iqgD***$^{L1-}$ **cre**). Introns are shown in gray, and the bsr cassette in purple. Arrowheads indicate some of the primers used to analyze the *iqgD* loci. Primers labeled in green were used to sequence the *iqgD*$^{L2}$ locus, while magenta primers were used to confirm a double knockout. The primers displayed below the gene were used to detect the presence of an alternative locus. (F) Schematic representation of the *iqgD* gene at the known locus on chromosome 2 before (***iqgD***$^{L1}$) and after (***iqgD***$^{L1-}$) homologous recombination, the *iqgD* at unknown genomic position before (***iqgD***$^{L2}$) and after (***iqgD***$^{L2-}$) homologous recombination; and *iqgD*$^{L1-}$ after excision of the bsr cassette with Cre recombinase (***iqgD***$^{L1-}$ **cre**). Introns are shown in gray, and the bsr cassette in purple. Arrowheads indicate some primers used to analyze the *iqgD* loci. Primers labeled in green were used to sequence the *iqgD*$^{L2}$ locus, while magenta primers were used to confirm a double knockout. The primers below the gene were used to detect the presence of an alternative locus. (G) Excision of the bsr cassette from *iqgD*$^{L1-}$ (I) and inactivation of *iqgD*$^{L2}$ (II) were confirmed by PCR. (I) Two PCRs were performed to detect the excision of the bsr cassette. One PCR was performed with a primer pair that yielded PCR products of different sizes for wt (2042 bp), *iqgD*$^{L1-}$ (3188 bp, barely visible) and *iqgD*$^{L1-}$ cre (1768 bp) cells (*upper gel*). The other PCR was performed with a primer pair that yielded PCR products of different sizes for wt (1214 bp), *iqgD*$^{L1-}$ (2360 bp) and *iqgD*$^{L1-}$ cre (940 bp) cells (*lower gel*). These PCR products were amplified from the first locus. A PCR-product of 1642 bp (with the 1st primer pair) and 814 bp (with the 2nd primer pair) of *iqgD*$^{L2}$ should be obtained in all cells. However, the other product was not obtained in sufficient quantity to be visible on the gel. (II) The successful generation of *iqgD*$^{L1-/L2-}$ cells was confirmed by two PCRs. One PCR was performed with a forward primer complementary to the gene sequence upstream of the 5' homology region and a reverse primer complementary to a bsr cassette, yielding a PCR product of 1181 bp (*left of the DNA standard on the gel*). The other PCR was performed with a forward primer complementary to the bsr cassette and another primer complementary to the gene sequence downstream of the 3' homology region and yielded 1840 bp (*right of the DNA standard on the gel*). Both PCR products can only be amplified from inactivated *iqgD*$^{L2}$ in the *iqgD*$^{L1-}$ cre background. The integration of the bsr cassette into *iqgD*$^{L2}$ was also confirmed by sequencing the 1840 bp product. The expression of IqgD in (C) was compared using Unpaired t-test with Welch's correction. The significance level was set at 5%. **** P < 0.0001.

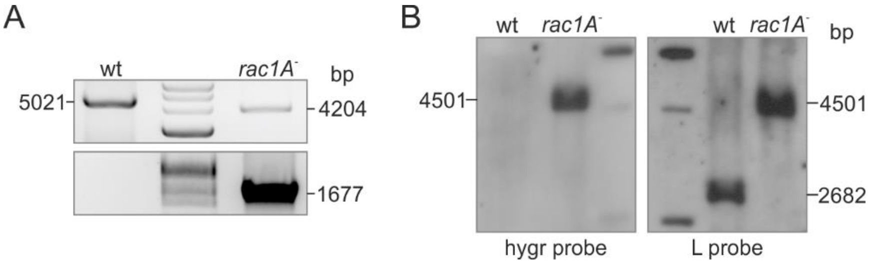


**Figure S7. Inactivation of the *rac1A* gene by homologous recombination was confirmed by (A) PCR and (B) Southern blot.** (A) Two PCRs were performed to detect the integration of a hygromycin resistance (hygr) cassette at the target locus. One PCR was performed with a primer pair that yielded PCR products of different sizes for wild-type (wt, 5021 bp) and *rac1A*$^-$ (4204 bp) cells (*upper gel*). The other PCR was performed with a primer complementary to a hygr cassette, while the other primer is complementary to the gene sequence upstream of the 5' homology region. Therefore, the PCR product (1677 bp) can only be obtained from *rac1A*$^-$ clone (*lower gel*). (B) Southern blot was performed with two probes, one complementary to the hygr cassette (hygr probe, *left membrane*) and the other complementary to the gene sequence upstream of the 5' homology region (L probe, *right membrane*). The hygr probe detects a band only in the *rac1A*$^-$ cells (4501 bp), while the L probe hybridizes with bands of different lengths in wild-type (2682 bp) and *rac1A*$^-$ (4501 bp) cells.

**Supporting Table**

**Table S1.** Sequences of oligonucleotides used in this work.

| Usage | Primer | Sequence 5' → 3' |
|---|---|---|
| ***Expression in D. discoideum cells*** | | |
| pDM304_YFP-IqgD | IqgD_N-term-F1 | TACAAATCCGGACTCAGATCTATGACATATACTGATAGTCGAT |
| | IqgD_N-term-R1 | ATTTGGATCCAAACCATTTGGTACACCTT |
| | IqgD_C-term-F1 | CAAATGGTTTGGATCCAAATAAAATGAAAA |
| | IqgD_C-term-R1 | TAATTTATTTATTTAACTAGTTTATAAACTAAATAATTTCATTAAT |
| pDM304_IqgD-YFP | IqgD_N-term-F2 | TAAAAAATAAAAATCAGATCTAAAAAATGACATATACTGATAGTCG |
| | IqgD_N-term-R2 | ATTTGGATCCAAACCATTTGGTACACCTTTAACTG |
| | IqgD_C-term-F2 | CAAATGGTTTGGATCCAAATAAAATGAAAA |
| | IqgD_C-term-R2 | TTCACCTTTACTACTACTAGTTAAACTAAATAATTTCATTAATAAA |
| pDM304_IqgD | IqgD_N-term-F2 | TAAAAAATAAAAATCAGATCTAAAAAATGACATATACTGATAGTCG |
| | IqgD_N-term-R2 | ATTTGGATCCAAACCATTTGGTACACCTTTAACTG |
| | IqgD_C-term-F1 | CAAATGGTTTGGATCCAAATAAAATGAAAA |
| | IqgD_C-term-R1 | TAATTTATTTATTTAACTAGTTTATAAACTAAATAATTTCATTAAT |
| pDM304_YFP-IqgD_ΔCHD | IqgD_N-term-F1 | TACAAATCCGGACTCAGATCTATGACATATACTGATAGTCGAT |
| | IqgD52-R2 | TTCTATGCTTTGTATGAACACTACCATCCTCAGCA |
| | IqgD325-F1 | TGTTCATACAAAGCATAGAACTGATCAA |
| | IqgD_C-term-R1 | TAATTTATTTATTTAACTAGTTTATAAACTAAATAATTTCATTAAT |
| pDM304_YFP-IqgD_ΔGRD | IqgD_N-term-F1 | TACAAATCCGGACTCAGATCTATGACATATACTGATAGTCGAT |
| | IqgD661-R1 | CTGTAACATATGGATCCAAACCATTTGGTACACCT |
| | IqgD1075-F1 | TTTGGATCCATATGTTACAGCCGCAAGAC |
| | IqgD_C-term-R1 | TAATTTATTTATTTAACTAGTTTATAAACTAAATAATTTCATTAAT |
| pDM304_YFP-IqgD_ΔRGCt | IqgD_N-term-F1 | TACAAATCCGGACTCAGATCTATGACATATACTGATAGTCGAT |
| | IqgD_N-term-R2 | ATTTGGATCCAAACCATTTGGTACACCTTTAACTG |
| | IqgD1291-F1 | CAAATGGTTTGGATCCAAATAAAATGAAAAACT |
| | IqgD1291-R1 | TAATTTATTTATTTAACTAGTTTAAATTGTTGATTTATCTTTTGGT |
| pDM330_DPAKa_GBD-mRFP | DPAKa_GBD-BglII-F1 | ATTAGATCTAAAAAATGGAGAGAGAAGATAAGAAAAAAG |
| | DPAKa_GBD-SpeI-R1 | AAGACTAGTCAATGGAATGGATGGTGC |
| pDM1208_IqgD | IqgD_N_pDM1208_BglII-F1 | GGTTCAGGAGGTAGTAGATCTATGACATATACTGATAGTCGAT |
| | IqgD_N-term-R1 | ATTTGGATCCAAACCATTTGGTACACCTT |
| | IqgD_C-term-F1 | CAAATGGTTTGGATCCAAATAAAATGAAAA |
| | IqgD_C-term-R1 | TAATTTATTTATTTAACTAGTTTATAAACTAAATAATTTCATTAAT |

| | | |
|---|---|---|
| pDM304_YFP-Lifeact | Lifeact_C-BglII-F1<br>Lifeact_C-SpeI-R1 | GATCTATGGGTGTCGCTGACCTGATAAAGAAGTTTGAAAGCATCTCCAAGGAAGAGTAAA<br>CTAGTTTACTCTTCCTTGGAGATGCTTTCAAACTTCTTTATCAGGTCAGCGACACCCATA |
| ***Yeast two-hybrid assay*** | | |
| pGBKT7_Rac1B(wt)_ΔCAAX | Rac1B_EcoRI-F1<br>Rac1B_dCAAX_BamHI-R1 | ATTAAGAATTCATGCAAGCAATTAAATGTGTAG<br>TAAGGATCCTTAACCTTTTGAAGATTTTGG |
| Rac1B(wt) to Rac1B(G12V) mutagenesis | Rac1B_G12V_mut-F1<br>Rac1B_G12V_mut-R1 | TGTTGGTGATGTTGCAGTTGGTA<br>ACTACACATTTAATTGCTTGC |
| Rac1B(wt) to Rac1B(T17N) mutagenesis | Rac1B_T17N_mut-F1<br>Rac1B_T17N_mut-R1 | AGTTGGTAAAAATTGTCTTTTAATTTCATATACAACCAATG<br>GCACCATCACCAACAACTAC |
| Rac1C(T17N) to Rac1C(wt) mutagenesis | Rac1C_N17T_mut-F1<br>Rac1C_N17T_mut-R1 | GGTTGGTAAAACATGTCTTTTAATTTCTTATACAACTAAC<br>GCACCATCACCTACAACTAC |
| Rac1C(wt) to Rac1C(G12V) mutagenesis | Rac1C_G12V_mut-F1<br>Rac1C_G12V_mut-R1 | TGTAGGTGATGTTGCGGTTGGTA<br>ACTACACATTTAATTGCTTGCATG |
| RacB(G12V) to RacB(wt) mutagenesis | RacB_V12G-mut-F1<br>RacB_V12G-mut-R1 | AGTAGGTGATGGTGCTGTTGGTA<br>ACCACCAATTTAATTGATTGCATG |
| RacB(wt) to RacB(Q61L) mutagenesis | RacB_Q61L_mut-F1<br>RacB_Q61L_mut-R1 | TACTGCTGGTCTAGAGGATTATG<br>TCCCAAAGACCTAATGAAAC |
| RacB(wt) to RacB(T17N) mutagenesis | RacB_T17N_mut-F1<br>RacB_T17N_mut-R1 | TGTTGGTAAAAATTGTTTATTAATTTCTTATAC<br>GCACCATCACCTACTACC |
| RacG(G12V) to RacG(wt) mutagenesis | RacG_V12G-mut-F1<br>RacG_V12G-mut-R1 | TGTTGGCGAAGGTGGAATTGGTA<br>ACACAAACTTTAATACTTTTCATGAATTCG |
| RacG(wt) to RacG(T17N) mutagenesis | RacG_T17N_mut-F1<br>RacG_T17N_mut-R1 | AATTGGTAAAAATTCAATGTTATTAAGTTATACATCAAATTCAATTTCAAATG<br>CCACCTTCGCCAACAACA |
| pGADT7_IqgD | N-IqgD_BglII-F1<br>N-IqgD_BamHI-R1 | ATTAGATCTCAATGACATATACTGATAGTCG<br>ATTTGGATCCAAACCATTTGGTAC |
| pGADT7_IqgD_GRD | GST-GRD_IqgD_BamHI-F1<br>Y2H-GRD_IqgD_SacI-R1 | ATTGGATCCAATAAAATGAAAAACTATC<br>AATGAGCTCTTATTTACCTTTATCCAAT |
| pGBKT7_IqgD | N-IqgD_BglII-F1<br>N-IqgD_BamHI-R1<br>C-IqgD_BamHI-F1<br>C-IqgD_SalI-R1 | ATTAGATCTCAATGACATATACTGATAGTCG<br>ATTTGGATCCAAACCATTTGGTAC<br>CCAAATGGTTTGGATCCAAATAAAATGA<br>ATTGTCGACTTATAAACTAAATAATTTCATTAATAAATG |
| ***BiFC assay*** | | |
| pDM304_VC-IqgD | N-IqgD-pDM304_VC-F1<br>IqgD_N-term-R1<br>IqgD_C-term-F1<br>C-IqgD-pDM304_VC-R1 | GGTTCTGGTTCAGGTAGATCTATGACATATACTGATAGTCGAT<br>ATTTGGATCCAAACCATTTGGTACACCTT<br>CAAATGGTTTGGATCCAAATAAAATGAAAA<br>TAATTTATTTATTTAACTAGTTTATAAACTAAATAATTTCATTAAT |
| pDM304_VC-IqgD_GRD | GST-GRD_IqgD_BamHI-F1<br>pDM304_VC-GRD_IqgD_SpeI-R1 | ATTGGATCCAATAAAATGAAAAACTATC<br>AATACTAGTTTATTTACCTTTATCCAAT |
| pDM304_VC-IqgD_ΔGRD | N-IqgD-pDM304_VC-F1<br>IqgD661-R1<br>IqgD1075-F1 | GGTTCTGGTTCAGGTAGATCTATGACATATACTGATAGTCGAT<br>CTGTAACATATGGATCCAAACCATTTGGTACACCT<br>TTTGGATCCATATGTTACAGCCGCAAGAC |

| | | |
|---|---|---|
| | IqgD_C-term-R1 | CAAATGGTTTGGATCCAAATAAAATGAAAA |
| pDM304_VC-IqgD_ΔRGCt | N-IqgD-pDM304_VC-F1 | GGTTCTGGTTCAGGTAGATCTATGACATATACTGATAGTCGAT |
| | IqgD_N-term-R1 | ATTTGGATCCAAACCATTTGGTACACCTT |
| | IqgD1291-F1 | CAAATGGTTTGGATCCAAATAAAATGAAAAACT |
| | IqgD1291-R1 | TAATTTATTTATTTAACTAGTTTAAATTGTTGATTTATCTTTTGGT |
| pDM304_VC-IqgD_311-1385 | IqgD_311_pDM304_VC-F1 | TGGTTCAGGTAGATCTTCAGAGGCTGCAATTGCA |
| | IqgD_N-term-R1 | ATTTGGATCCAAACCATTTGGTACACCTT |
| | IqgD_C-term-F1 | CAAATGGTTTGGATCCAAATAAAATGAAAA |
| | C-IqgD-pDM304_VC-R1 | TAATTTATTTATTTAACTAGTTTATAAACTAAATAATTTCATTAAT |
| pDM344_VN-Rac1C(G12V) /(T17N) | Rac1C_BamHI-F1 | ATTGGATCCATGCAAGCAATTAAATGTG |
| | Rac1C_SpeI-R1 | AATACTAGTTTACAAGATATTACATCCACTTTTAC |
| Rac1A(wt) to Rac1A(G12V) mutagenesis | Rac1A_G12V_fl_BglII-mut-F1 | TGTCGGTGATGTTGCTGTAGGTA |
| | Rac1A_G12V_fl_SpeI-mut-R1 | ACGACACATTTAATTGCTTGC |
| Rac1C(wt) to Rac1C(G12) mutagenesis | Rac1C_G12V_mut-F1 | TGTAGGTGATGTTGCGGTTGGTA |
| | Rac1C_G12V_mut-R1 | ACTACACATTTAATTGCTTGCATG |
| Rac1C(wt) to Rac1C(T17N) mutagenesis | Rac1C_T17N-mut-F1 | GGTTGGTAAAAATTGTCTTTTAATTTCTTATACAACTAAC |
| | Rac1C_N17T_mut-R1 | GCACCATCACCTACAACTAC |
| Rac1C(wt) to Rac1C(Q61L) mutagenesis | Rac1C_Q61L_mut-F1 | TACTGCTGGTCTAGAAGATTATG |
| | Rac1C_Q61L_mut-R1 | TCCCACAAGCCAAGATTAATTG |
| ***Production of GST-tagged IqgD and Rac1A for purification from bacterial cells*** | | |
| pGEX-6P-1_IqgD_1-575 | GST-CH_IqgD_BamHI-F1 | ATTGGATCCATGACATATACTGATAGTC |
| | GST-N_term-IqgD_SalI-R1 | AATGTCGACTTAAATTTTATATTCTTTATTGT |
| pGEX-6P-1_Rac1A(Q61L) | GST-Rac1A_wt_BamHI-F1 | ATTGGATCCATGCAAGCAATTAAATGTG |
| | GST-Rac1A_wt_SalI-R1 | AATGTCGACTTATAAAATGTTGCAACCAC |
| pGEX-6P-1_Rac1A(T17N) | GST-Rac1A_wt_BamHI-F1 | ATTGGATCCATGCAAGCAATTAAATGTG |
| | GST-Rac1A_wt_SalI-R1 | AATGTCGACTTATAAAATGTTGCAACCAC |
| ***Production of StrepII-tagged IqgD proteins for purification from insect cells*** | | |
| pCoofy51_IqgD_1-592 | IqgD_pCoofy_N-F1 | TTCTGTTCCAGGGGCCCATGACATATACTGATAGTCGATATAAT |
| | IqgD_592_pCoofy-R1 | TACCGCATGCCTCGAGTTAATTATCAGTGAGTGAGGTATC |
| pCoofy51_IqgD_311-1385 | IqgD_311_pCoofy-F1 | TTCCAGGGGCCCATGTCAGAGGCTGCAATTGCA |
| | IqgD_N-term-R1 | ATTTGGATCCAAACCATTTGGTACACCTT |
| | IqgD_C-term-F1 | CAAATGGTTTGGATCCAAATAAAATGAAAA |
| | IqgD_pCoofy_C-R1 | CTTGGTACCGCATGCCTCGAGTTATAAACTAAATAATTTCATTAAT |
| ***Production of iqgD$^{-}$ cells*** | | |
| *iqgD*-L1-3’-fr | IqgD_3'-PstI-F1 | ATTCTGCAGGATTGATAGATCAAGAGGATG |
| | IqgD_3'-BamHI-R1 | ATTGGATCCCACTATCTTGAATATCCTCAC |
| PCR for *iqgD*$^{L1-}$ selection | IqgD-gDNA-F1 | ATTCCCATTTATTGCTATTATCAAATTTC |
| | IqgD-gDNA-R1 | AATACACCAATCTTTTCACATGCC |

| | | |
|---|---|---|
| | Bsr-UP<br>IqgD-D1 | AGTATTCGAGTGGTAAGTCCTTG<br>ACACCAATCTTTTCACATGCC |
| PCR for sequencing $iqgD^{L2}$ in $iqgD^{L1-}$ cells | IqgD-HindIII-F1<br>iqgD_probe-rev1<br><br>IqgD_seq-for1-L2<br>IqgD_seq-rev2-L2 | ATTAAGCTTATGACATATACTGATAGTC<br>CTTCTAGACGTAAGGCTTCTTC<br><br>GCTTATTCAAAATTTATAAC<br>CATCTTGAATTGAAAATAGATAC |
| PCR for $iqgD^{L1-}$ creed selection | IqgD-HindIII-F1<br>iqgD_probe-rev1<br><br>IqgD_5'-U1<br>IqgD-D1 | ATTAAGCTTATGACATATACTGATAGTC<br>CTTCTAGACGTAAGGCTTCTTC<br><br>GAGGTAGCAAGAAAATGGAC<br>ACACCAATCTTTTCACATGCC |
| *iqgD*-L2-3'-fr | IqgD-L2-3'-PstI-F1<br>IqgD-L2-3'-BamHI-R1 | ATTCTGCAGCGTGAACGTATAGAAAGGGAG<br>ATTGGATCCCAAATTCTTCACGTTCAATACG |
| *iqgD*-L2-5'-fr | IqgD-L2-5'-SalI-F1<br>IqgD-L2-5'-HindIII-R1 | ATTGTCGACAGAGGTAGCAAGAAAATGGAC<br>TCCAAGCTTCCTCTGCTGAATGTTCATCTG |
| PCR for $iqgD^{L1-/L2-}$ confirmation | IqgD-HindIII-F1<br>Bsr<br><br>Bsr-UP<br>IqgD_endBamHI-R1 | ATTAAGCTTATGACATATACTGATAGTC<br>CAGTTACTCGTCCTATATACG<br><br>AGTATTCGAGTGGTAAGTCCTTG<br>ATTTTATTTGGATCCAAACCATTTG |
| PCR for sequencing $iqgD^{L2-}$ | Bsr-UP<br>IqgD_endBamHI-R1 | AGTATTCGAGTGGTAAGTCCTTG<br>ATTTTATTTGGATCCAAACCATTTG |
| bsr probe for Southern | Bsr_Southern-UP2<br>Bsr_Southern-DP | TTCTTATTTCTTAAAACAAATAAATTAA<br>TTAATTTCGGGTATATTTGAGTG |
| R probe for Southern | iqgD_probe-for1<br>iqgD_probe-rev1 | CGTGAACGTATAGAAAGGGAG<br>CTTCTAGACGTAAGGCTTCTTC |
| ***qPCR and end-point PCR for the presence of*** $iqgD^{L2}$ | | |
| *gpdA* | gpdA_qPCR-F1<br>gpdA_qPCR-R1 | TTTTGGTCGTATCGGTCGTC<br>TACCTTTGAAACGACCGTGG |
| *iqgD* | IqgD_RT-UP1<br>IqgD_RT-DP1<br><br>IqgD_RT-DP2<br>IqgD_RT-DP3<br>IqgD_RT-UP4<br>IqgD_RT-DP4 | GGATGGTAGTGTTCATACATTC<br>AGTTGATGGGTCAATTGGAAC<br><br>CCAAATTACACCTAATACTAAATAGG<br>CAACTTCTTTAACTGAATCTTCTTC<br>TGAAGAAGAAAAAGTTGCTTATTC<br>TAAAACACCATCTGTACAATTTAC |
| ***Production of*** $rac1A^{-}$ ***cells*** | | |
| PCR for $rac1A^{-}$ selection | 5'check pDM1490-UP<br>3'check pDM1490-DP<br><br>hygR_for-seq3<br>3'check pDM1490-DP | CATCAGTTAATGGTTGAATTGTTAATGG<br>GAAGATGAAACCTTTATACCAAAACTTG<br><br>ACTCGTCCGGATCGGGAG<br>GAAGATGAAACCTTTATACCAAAACTTG |
| hygr probe for Southern | hygR_probe-F1<br>hygR_probe-R1 | GCTTTCAGCTTCGATGTAGGAG<br>CCAGTCAATGACCGCTGTTATG |
| L probe for Southern | 1A-Sou-ClaI-UP<br>1A-Sou-ClaI-DP | CAACATAATTTCAATTAATTCTCTC<br>ATCAAAGAATAAAAGATTTACC |

**Legends for Movies S1-S17**

**Movie S1**: Time-lapse recording of a randomly moving vegetative wild-type cell expressing IqgD-YFP, acquired by confocal microscopy. Refers to Fig. 1A. Scale bar: 5 µm.

**Movie S2**: Time-lapse recording of a randomly moving vegetative wild-type cell expressing YFP-IqgD, acquired by confocal microscopy. Refers to Fig. 1B. Scale bar: 5 µm.

**Movie S3**: Time-lapse recording of a vegetative wild-type cell expressing YFP-IqgD during ingestion of a TRITC-labeled yeast particle, acquired by confocal microscopy. Refers to Fig. 1C. Scale bar: 5 µm.

**Movie S4**: Time-lapse recording of a vegetative wild-type cell expressing YFP-IqgD during cytokinesis, acquired by confocal microscopy. Refers to Fig. 1D. Scale bar: 10 µm.

**Movie S5**: Time-lapse recording of a vegetative wild-type cell coexpressing IqgD-YFP and Lifeact-mRFP during phagocytosis of yeast particle, acquired by confocal microscopy. Refers to Fig. S1A. Scale bar: 5 µm.

**Movie S6**: Time-lapse recording of a vegetative wild-type cell coexpressing YFP-IqgD and Lifeact-mRFP during random movement and macropinocytosis, acquired by confocal microscopy. Refers to Fig. S1B. Scale bar: 5 µm.

**Movie S7**: Time-lapse recording of a vegetative *iqgD*$^{L1-}$ cell expressing YFP-IqgD_ΔCHD during random movement and macropinocytosis, acquired by confocal microscopy. Refers to Fig. 2B. Scale bar: 5 µm.

**Movie S8**: Time-lapse recording of a vegetative wild-type cell coexpressing YFP-IqgD and DPAKa_GBD-mRFP during random movement and macropinocytosis, acquired by confocal microscopy. Refers to Fig. 3B. Scale bar: 5 µm.

**Movie S9**: Time-lapse recording of a vegetative *iqgD*$^{L1-}$ cell expressing VC-IqgD and VN-Rac1A(wt) during random movement and macropinocytosis, acquired by confocal microscopy. Refers to the upper panel of Fig. 3D. Scale bar: 5 µm.

**Movie S10**: Time-lapse recording of a vegetative *iqgD*$^{L1-}$ cell expressing VC-IqgD and VN-Rac1B(wt) during random movement and macropinocytosis, acquired by confocal microscopy. Refers to the middle panel of Fig. 3D. Scale bar: 5 µm.

**Movie S11**: Time-lapse recording of a vegetative *iqgD*$^{L1-}$ cell expressing VC-IqgD and VN-Rac1C(wt) during random movement and macropinocytosis, acquired by confocal microscopy. Refers to the lower panel of Fig. 3D. Scale bar: 5 µm.

**Movie S12**: Time-lapse recording of a vegetative *iqgD*$^{L1-}$ cell expressing YFP-IqgD_ΔGRD during random movement and macropinocytosis, acquired by confocal microscopy. Refers to the upper panel of Fig. 4B. Scale bar: 5 µm.

**Movie S13**: Time-lapse recording of a vegetative *iqgD*$^{L1-}$ cell expressing YFP-IqgD_ΔRGCt during random movement and macropinocytosis, acquired by confocal microscopy. Refers to the lower panel of Fig. 4B. Scale bar: 5 µm.

**Movie S14**: Time-lapse recording of a vegetative *iqgD*$^{L1-}$ cell expressing YFP-IqgD and Lifeact-mRFP during random movement, acquired by TIRF microscopy. Scale bar: 5 µm.

**Movie S15**: Time-lapse recording of a vegetative wild-type cell expressing YFP-Lifeact, acquired by confocal microscopy. The focus was adjusted so that fat droplets at the glass surface were sharply visible. Refers to Fig. 5K. Scale bar: 5 µm.

**Movie S16**: Time-lapse recording of a vegetative wild-type cell expressing YFP-IqgD, acquired by confocal microscopy. The focus was adjusted so that fat droplets at the glass surface were sharply visible. Refers to Fig. 5K. Scale bar: 5 µm.

**Movie S17**: Time-lapse recording of a vegetative wild-type cell expressing DPAKa_GBD-YFP, acquired by confocal microscopy. The focus was adjusted so that fat droplets at the glass surface were sharply visible. Refers to Fig. 5K. Scale bar: 5 µm.

**Supporting Information References**
